# Intrinsically linked lineage specificity of transposable elements, lncRNA genes, and transcriptional regulation

**DOI:** 10.1101/2024.03.04.583292

**Authors:** Jie Lin, Yijin Wu, Junxiang Zeng, Wei Xiong, Sha He, Pierre Pontarotti, Hao Zhu

## Abstract

Unlike protein-coding genes, many mammalian lncRNAs are lineage-specific (LS), but which human and mouse reference lncRNAs are lineage-specific, how they are generated, and whether their regulatory elements are lineage-specific remain unclear. To resolve their origins, properties, and impacts, we systematically classified primate-, simian-, rodent-, human-, and mouse-specific lncRNAs in GENCODE-annotated human and mouse genomes using structure-aware covariance modeling. We predicted their DNA-binding domains (DBDs) and genomic binding sites (DBSs) and traced the origins of embedded transposable elements (TEs). LS TEs contribute to over 84% (human) and 98% (mouse) of LS lncRNAs, 61% and 95% of their DBDs, and 46% and 73% of their DBSs. Analyses of multi-omics data and a developmental process suggest that LS TE-linked lineage-specificity substantially rewires transcriptional regulation and that many evolutionary novelties may be structurally predisposed. These findings bridge the gap between human diseases and mouse models, providing valuable resources for identifying human-specific therapeutic lncRNA targets.

## 1. Introduction

Despite extensive conservation of protein-coding genes (PCGs) across mammals, substantial differences in development, physiology, and phenotype persist. Uncovering the molecular mechanisms underlying the gaps between genotypes and phenotypes and between human diseases and their corresponding animal models remains a long-standing challenge. While these gaps are determined by lineage-specific (LS) genes and LS regulatory elements (King and Wilson, 1975), three questions remain unsolved: which genes and regulatory elements are lineage-specific, what determines their lineage specificity, and whether LS genes and LS regulatory elements are co-opted.

Studies have reported that the vast majority of mammalian lncRNAs are lineage- or species-specific (Hezroni et al., 2015; Necsulea et al., 2014; Sarropoulos et al., 2019; Washietl et al., 2014). However, since these studies relied on *de novo* transcript assembly from organ-derived RNA-seq data across multiple species rather than systematically categorizing the evolutionary origins of standard, validated reference lncRNAs, it remains unclear which GENCODE-annotated lncRNAs in humans and mice are lineage-specific. Further, since these studies used sequence-only alignment tools (i.e., BLAST and Multiz) to infer noncoding orthology, their results are prone to false negatives due to the rapid sequence turnover of non-coding transcripts.

Transposable elements (TEs) critically contribute to the birth of new genes (Cosby et al., 2021; Ruiz-Orera et al., 2015) and new regulatory elements (e.g., transcription factor DNA binding sites (TFBS)) (Bourque et al., 2018; Fueyo et al., 2022; Wells and Feschotte, 2020). However, previous studies did not systematically examine whether TEs contribute to the regulatory elements of LS lncRNAs. An important function of lncRNAs is transcriptional regulation. Many lncRNAs can localize to specific genomic sites by forming ncRNA-DNA triplexes and subsequently recruiting epigenetic modifiers to these sites (Chu et al., 2011; Engreitz et al., 2016; Yap et al., 2010), but whether TEs substantially contribute to ncRNA-DNA interaction remains uncharacterized.

A substantial fraction of GENCODE-annotated human and mouse lncRNAs are lineage-specific (Derrien et al., 2012; Yue et al., 2014); species-specific lncRNAs can alter gene expression in conserved tissues and cells (Breschi et al., 2017; Hodge et al., 2019); and human-specific (HS) lncRNAs may have greatly promoted human evolution (Lin et al., 2026b). However, a systematic evolutionary classification of GENCODE-annotated lncRNAs remains unreported. Mapping the lineage specificity of these reference lncRNAs in the focal genomes of humans and mice is essential because diverse mouse models are used to study human diseases, abundant transcriptomic data enable comparative lncRNA functional analysis, and focusing on reference annotations allows LS lncRNAs to be directly mapped to clinical and downstream disease datasets. In contrast, unannotated, *de novo*-assembled LS lncRNA transcripts cannot be easily integrated with established clinical databases.

We postulate that LS TEs are structurally linked to the emergence of both LS lncRNAs and their regulatory elements. To test this hypothesis, this study systematically classifies primate-specific (PS), simian-specific (SS), rodent-specific (RS), human-specific (HS), and mouse-specific (MS) lncRNAs within the GENCODE reference annotations of humans and mice. Our methods are distinct from prior studies. (a) Structure-Aware Homology Profiling: We searched for orthologs of human and mouse lncRNA exons across mammalian genomes using the *Infernal* program, which executes covariance model-based homology searches that incorporate both nucleotide sequence and RNA secondary structure information (Nawrocki and Eddy, 2013). (b) Triplex-Mediated Domain Prediction: We predicted DNA-binding domains (DBDs) in LS lncRNAs and their DNA-binding sites (DBSs) in promoter regions of all Ensembl-annotated transcripts. Triplex formation between DBDs and DBSs follows strict base-pairing rules (Abu Almakarem et al., 2012), making DBDs and DBSs computationally predictable and enabling systematic analysis of LS lncRNA functions (He et al., 2026; Lin et al., 2026b). (c) Tracing TE Origins: We traced the evolutionary origins of every TE overlapping LS lncRNAs or their DBDs and DBSs across 87 representative species spanning multiple lineages. By analyzing data at levels of genomic sequences, Gene Ontology (GO) categories, signaling pathways, and normal/diseased tissue transcriptomes, we demonstrate that LS TEs intrinsically link LS lncRNAs, their DBDs and DBSs, and lineage-specific transcriptional regulation. This link provides a highly flexible, low-pleiotropy mechanism for mammals to rapidly rewire transcriptional regulation (Kluger et al., 2004). The identified 4,864 and 4,125 LS lncRNAs in humans and mice provide a valuable resource for further computational and experimental investigations, including uncovering the molecular mechanisms underlying gaps between genotypes and phenotypes and between human diseases and their corresponding animal models, and identifying human- and disease-specific lncRNA targets.

## 2. Results

### 2.1 LS TEs contributed to LS lncRNAs and their regulatory elements

#### Identification of LS lncRNAs

*Using Infernal* to search for orthologs of exons of 13,562 GENCODE-annotated human lncRNAs (V18) across 16 mammalian genomes, we identified 66 HS lncRNAs (Lin et al., 2026a). Using the same method, we searched for orthologs of exons of 13,450 GENCODE-annotated mouse lncRNAs (M22) in *Mus caroli, Mus pahari*, rat, and rabbit and identified 212 MS lncRNAs (Supplementary Tables 1, 2). We then identified PS and SS lncRNAs in humans by examining the presence of orthologs of human lncRNA exons in three simians (chimpanzee, macaque, marmoset), three prosimians (tarsier, mouse lemur, tree shrew), and the mouse, and identified RS lncRNAs in mice by examining the presence of orthologs of mouse lncRNA exons in three rodents (*Mus caroli, Mus pahari*, rat) and rabbit. 4,864 LS (including HS) lncRNAs in humans and 4,125 LS (including MS) lncRNAs in mice were identified (Supplementary Tables 3, 4).

#### Contribution of LS TEs to LS lncRNAs

We examined TE contributions to HS and MS lncRNAs using RepeatMasker (http://repeatmasker.org/). 84% of HS lncRNAs and 98% of MS lncRNAs contain at least one exon with >30% of its length overlapping annotated TEs, and TE families in HS and MS lncRNAs are distinct. Compared with overall GENCODE-annotated human and mouse lncRNAs, TE coverage in HS and MS lncRNAs is significantly higher (one-sided Mann–Whitney test) (Fig. 1A–D).

**Figure 1.**
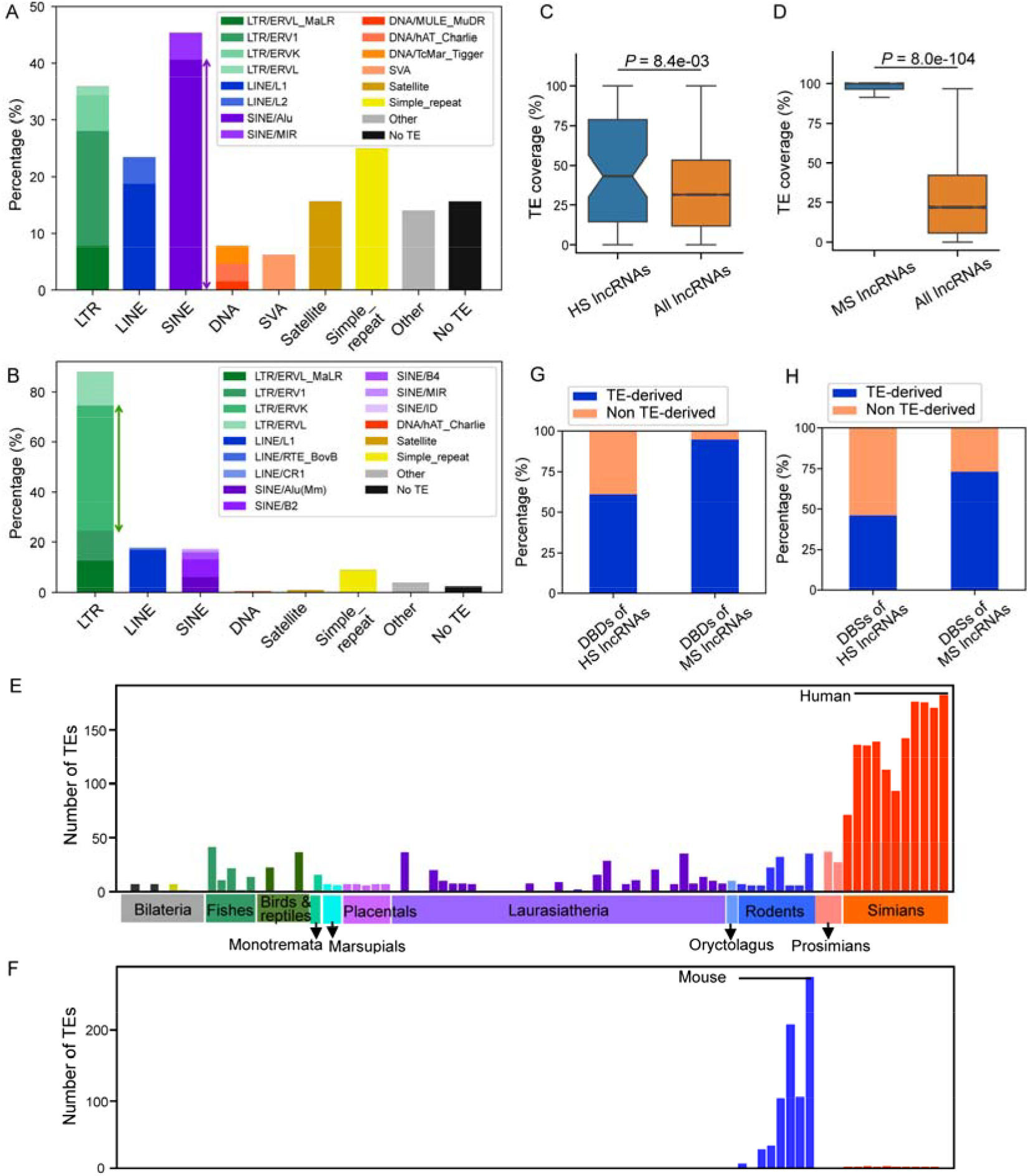
Simian/rodent TEs contribute sequences to HS/MS lncRNAs and their DBSs. (AB) Percentages of HS/MS lncRNAs with exons overlapping specific TE classes. (CD) TE coverage of HS/MS lncRNAs compared with TE coverage of entire GENCODE-annotated human/mouse lncRNAs. (EF) TEs in HS/MS lncRNAs are almost exclusively of simian/rodent origins. (GH) Percentages of HS/MS lncRNAs’ DBDs and DBSs that overlap with ≥ one TE by >30% of their length.

To examine whether these TEs are lineage-specific, we developed a phylogeny-based pipeline to search each TE across 87 representative species spanning major animal clades (Supplementary Figs. 1-3; Supplementary Table 5). Hits that have the top score in each species and also have normalized scores >0.4 were aligned using *MAFFT* (Katoh and Standley, 2013). Based on phylogenetic trees of hits reconstructed using *RAxML* (Stamatakis, 2014), 64% of TEs overlapping HS lncRNAs are restricted to simians, and 99% of TEs overlapping MS lncRNAs are restricted to rodents (Fig. 1EF; Supplementary Figs. 4, 5).

#### Contribution of LS TEs to DBDs and DBSs

We used the *LongTarget* program to predict DBDs within LS lncRNAs and the corresponding DBSs in promoter regions of Ensembl-annotated transcripts (179,128 in GRCh38.79 and 95,052 in GRCm38.83) (Lin et al., 2019). DBDs were predicted in 64 HS and 208 MS lncRNAs, with 102,651 and 566,581 DBSs (mean length = 149 bp and 147 bp) predicted in human and mouse transcripts (Supplementary Tables 6, 7). In humans, 61% of DBDs and 46% of DBSs overlap simian TEs by >30% of their length; in mice, 95% of DBDs and 73% of DBSs overlap rodent TEs by >30% of their length (Fig. 1GH).

#### Preferential interaction between LS lncRNAs and LS DBSs

Because ncRNA-DNA triplexes follow specific base-pairing rules, DBSs across species may share common sequence features, undermining the lineage specificity of transcriptional regulation. To address whether LS lncRNAs preferentially bind to LS DBSs, we analyzed spermatogenesis, a process intensively regulated by LS lncRNAs. Of the 4,864 and 4,125 LS lncRNAs in humans and mice, 599 PS/SS/HS lncRNAs and 1,196 RS/MS lncRNAs (having consistent gene symbols across Ensembl releases) were identified in a reported study (Murat et al., 2023) (Supplementary Table 8). We predicted their DBSs in 395 spermatogenesis markers (conserved since monotremes), and tested whether DBSs overlapping LS TEs show preferential binding to lncRNAs from the same lineage. DBSs of lncRNAs from the same lineage and overlapping TEs from the same lineage outnumber and/or have higher binding affinity than DBSs of lncRNAs from different lineages and overlapping TEs from different lineages (Supplementary Fig. 8; Supplementary Tables 9).

#### Evidence for transcriptional rewiring

We further examined whether LS lncRNAs regulate genes by targeting newly emerged or repurposed conserved genomic sequences. Of the 102,651 DBSs predicted in humans, 76,992 fall within human–mouse syntenic blocks, 44,525 have identifiable mouse counterparts, 33,947 fit both, and only 15,081 are both non-syntenic and lack mouse counterparts. These findings suggest transcriptional regulation arises from modifying the existing genomic architecture rather than exclusively from *de novo* sequence creation. The contribution of LS TEs to LS lncRNAs’ target PCGs also supports LS TE-mediated transcriptional rewiring (Supplementary Figs. 6, 7; Supplementary Tables 10, 11).

### 2.2 HS/MS lncRNAs’ target genes are enriched in distinct biological processes

First, we evaluated the functions of HS/MS lncRNAs by applying over-representation analysis (ORA) to their target genes. We ranked target genes by the number of DBSs, identified the top 10% as mostly regulated targets, and used the whole genome as background. *g:Profiler* and *PANTHER* reveal that HS lncRNA targets are significantly enriched for neurodevelopmental Gene Ontology (GO) terms, but MS lncRNA targets are enriched for broader developmental categories (Fig. 2A). Direct comparison of human top targets against mouse top targets indicates that the former and latter are enriched for neural development and immune-related GO terms and sensory perception GO terms, respectively (Fig. 2B).

**Figure 2.**
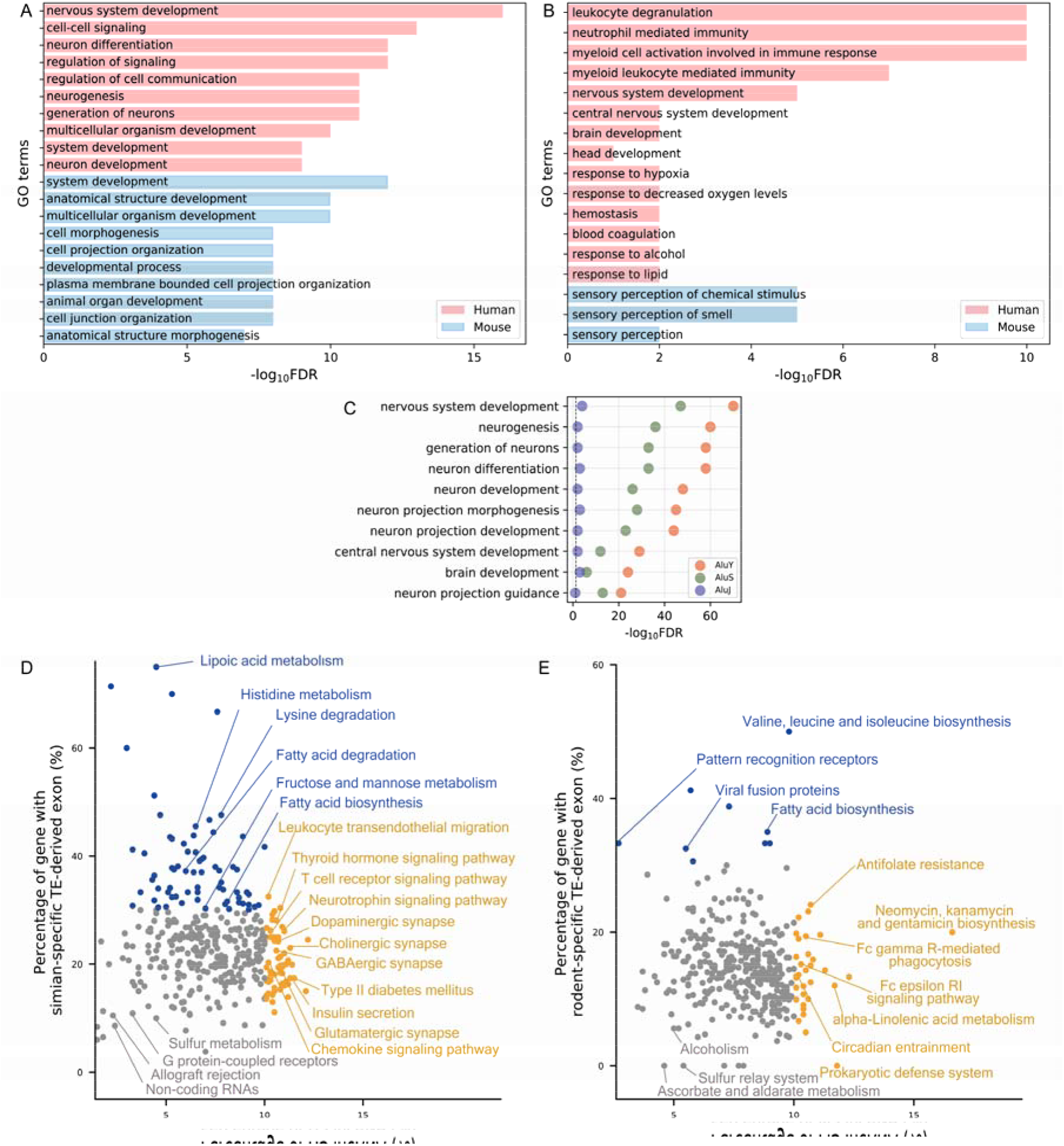
Functional enrichment and pathway distribution of HS/MS lncRNA target genes. (A) Top 10 enriched GO terms of HS lncRNAs’ top 10% target genes and MS lncRNAs’ top 10% target genes. (B) GO terms of HS lncRNAs’ top 10% target genes (foreground) and MS lncRNAs’ top 10% target genes (background). (C) GO enrichment of target genes of HS lncRNAs whose DBDs contain AluJ-, AluS-, and AluY-derived DBDs. (DE) Distribution of KEGG pathways according to the percentage of genes containing simian (D) or rodent (E) TE-derived exons (y-axis) and the percentage of genes containing DBSs from >10% HS/MS lncRNAs (x-axis). Pathways with >10% genes containing HS/MS DBSs are shown in orange; pathways with >30% genes containing simian/rodent TE-derived exons are shown in blue; other pathways are shown in grey.

Alu elements are short PS TEs and fall into three subfamilies based on age (AluJ, ∼65 MYA; AluS, ∼30 MYA; AluY, youngest) (Deininger, 2011; Larsen et al., 2018). Since Alu-derived sequences are present in the DBD of several HS lncRNAs, we examined whether Alu age correlates with functional enrichment of target genes. HS lncRNAs containing AluY-derived DBDs (n = 2) show the strongest enrichment for neurodevelopment-related GO terms, followed by AluS-derived (n = 7) and AluJ-derived (n = 2) groups (Fig. 2C). This pattern supports the link between LS TEs, LS lncRNAs, and lineage-specific transcriptional regulation.

Next, we examined the impact of HS/MS lncRNAs on molecular signaling. The Notch, Hippo, Wnt, and Hedgehog pathways contain important targets for anti-cancer drugs (Clara et al., 2020; Dey et al., 2020), and genes in these pathways frequently contain simian/rodent TE-derived exons and HS/MS lncRNA DBSs (Supplementary Figs. 9, 10). We then extended the analysis to all 382 KEGG pathways and found that 142 pathways had either a high percentage of genes containing simian/rodent TE-derived exons or a high percentage of genes containing DBSs of >10% HS and MS lncRNAs (Fig. 2DE).

### 2.3 HS/MS lncRNAs show species- and tissue-specific expression associations

Next, we examined the tissue specificity of HS/MS lncRNA-associated gene expression using the GTEx human data (54 tissues) and mouse transcriptomic data (17 tissues) (GTEx_Consortium., 2017; Schaum et al., 2020). We computed Spearman correlations between HS/MS lncRNAs and their target genes within each tissue. HS lncRNA-target gene pairs with significant expression correlation (|Spearman’s ρ| > 0.3, FDR < 0.05) are most prevalent in brain tissues; in contrast, MS lncRNA-target gene correlations are strongest in skin and bone tissues (Fig. 3AB). This result is consistent with the functional enrichment analysis and highlights high tissue specificity of transcriptional regulation by HS/MS lncRNAs.

**Figure 3.**
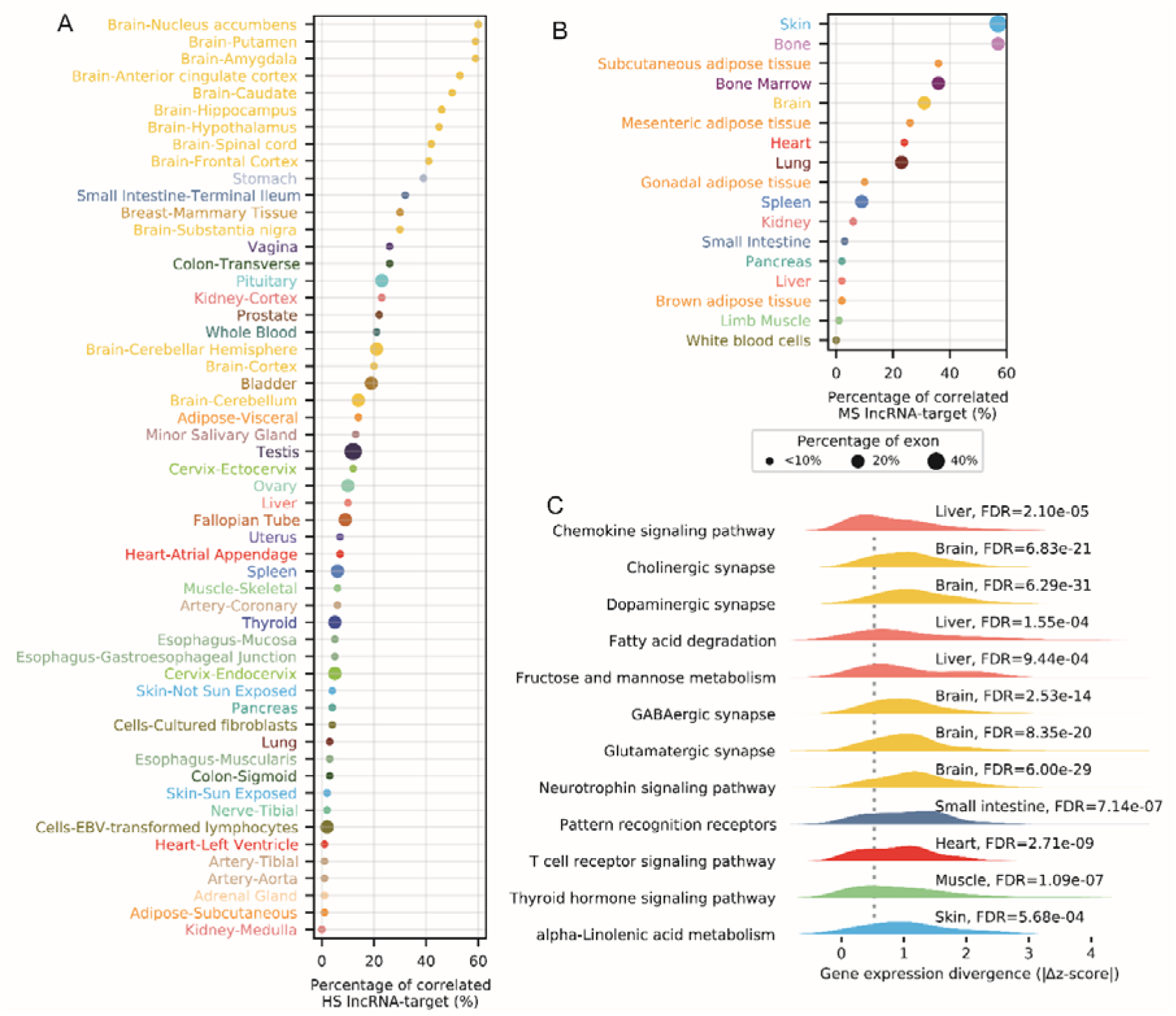
Impacts of simian/rodent TEs and HS/MS lncRNAs on gene expression and molecular signaling. (AB) Percentages of HS (A) and MS (B) lncRNA-target gene pairs showing significant expression correlation across human (54 GTEx tissues) and mouse (17 tissues) transcriptomes. Dot sizes indicate the percentage of target genes that contain simian/rodent TE-derived exons. (C) Density distributions of cross-species |Δz| values indicate that tissue-specific divergence in cross-species expression of one-to-one orthologous genes also exhibits pathway specificity. Dashed vertical lines indicate the median absolute z-score difference.

We also examined whether target genes’ TE-derived exons show tissue preference by normalizing TPM values of simian/rodent TE-derived exons into z-scores across tissues. Notably, simian TE-derived exons are preferentially expressed in the testis, whereas rodent TE-derived exons are preferentially expressed in the skin (Fig. 3AB).

To determine whether tissue-specific expression is associated with pathway-level divergence, we jointly analyzed one-to-one orthologous PCGs in (a) the 142 KEGG pathways identified above (Fig. 2DE) and (b) the 11 tissues with transcriptomic data available for both humans and mice. For each pathway in each tissue, we computed cross-species expression divergence as the absolute difference in z-scores between PCGs (Fig. 3C). Using pancreas as the reference (which shows a low proportion of correlated HS/MS lncRNA-target gene pairs in both species), 89% (126/142) of these KEGG pathways display significantly greater cross-species expression divergence in at least one tissue (one-sided Mann–Whitney test, FDR < 0.001) (Supplementary Table 12). In brain tissues, pathways with significant divergence include Cholinergic synapse, Dopaminergic synapse, GABAergic synapse, Glutamatergic synapse, and Neurotrophin signaling pathway. These results suggest that pathways enriched for species-specific TE-lncRNA associations frequently coincide with tissues exhibiting elevated cross-species expression divergence.

### 2.4 Transcriptional regulation by HS/MS lncRNAs in AD patients and mouse models

The cross-species differences in gene expression in normal tissues suggest that transcriptional regulation by HS/MS lncRNAs may cause substantial differences between human diseases and their mouse models. Alzheimer’s disease (AD) is characterized by extensive dysregulated genes and pathways (Guo et al., 2020; Hampel et al., 2021; Zhou et al., 2020). Many AD mouse models have been developed, but systematic analyses reveal that a substantial fraction of gene expression changes in mouse models differ from those observed in human patients (Supplementary Fig. 11) (Preuss et al., 2020; Wan et al., 2020). To examine whether HS/MS lncRNAs are associated with species-specific regulatory organization in AD, we analyzed RNA-seq datasets from AD patients (n

= 12) and a mouse model expressing humanized amyloid-β (n = 14) (Supplementary Table 13) (Baglietto-Vargas et al., 2021; Nativio et al., 2018). Using genes from AD-related KEGG pathways— particularly nervous and immune system pathways—as input (Benarroch, 2018; Hampel et al., 2018; Zott and Konnerth, 2023), we identified transcriptional regulatory modules with distinct species-specific features using the *eGRAM* program (Fig. 4). Several notable differences emerged. First, the number of regulatory modules differs substantially between humans and mice (26 vs 10). Second, individual HS lncRNAs are connected to more genes and more modules than MS lncRNAs. Third, *APP* and *MAPT*, which are major therapeutic targets in AD studies (Self and Holtzman, 2023), contain DBSs of both HS and MS lncRNAs; however, they are incorporated into regulatory modules only in humans. In contrast, *APOE*, a major late-onset AD risk gene (Knopman et al., 2021), contains strong DBSs exclusively from MS lncRNAs (Supplementary Table 14). Fourth, many modules contain gene pairs that show species-specific expression correlation (|Spearman’s ρ| > 0.5, FDR < 0.05 in one species, but |Spearman’s ρ| < 0.1, FDR > 0.05 in the other) (Fig. 4C), indicating differential co-regulation between human patients and mouse models. These differences provide a novel framework for uncovering transcriptional discrepancies between human patients and mouse models and resolving translational failures in AD drug development (Huang et al., 2020; Mehta et al., 2017; Zhang et al., 2023).

**Figure 4.**
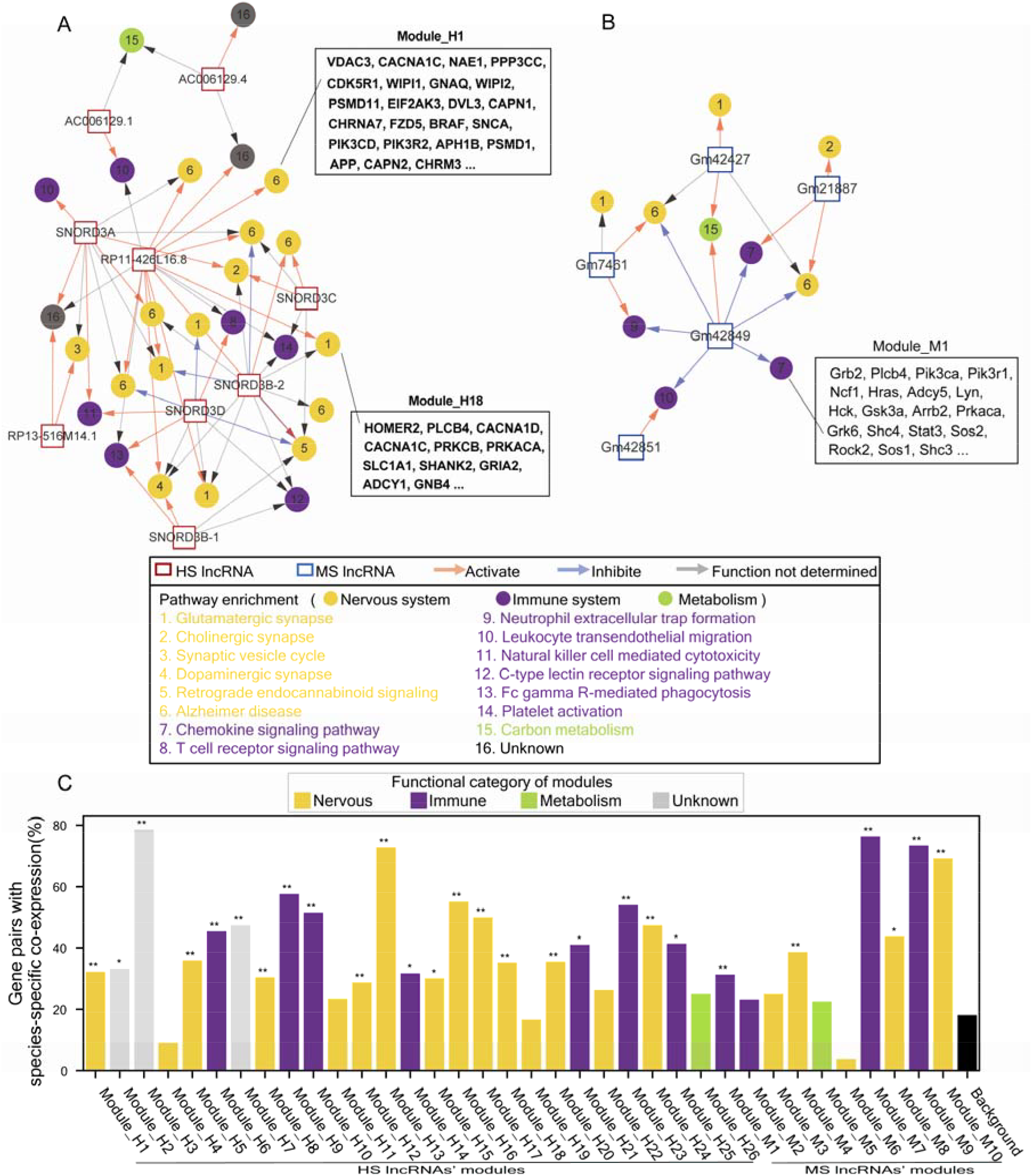
Species-specific transcriptional regulatory modules and networks of AD-related genes in humans and mice. (AB) Regulatory networks constructed from AD-related pathway genes and HS/MS lncRNAs in human patients (A) and mouse modules (B). Colored squares indicate lncRNAs. Colored circles indicate modules significantly enriched for genes in the numbered pathways (hypergeometric distribution test, FDR < 0.05). Pink, grey, and light grey arrows indicate activation, inhibition, and undetermined regulation. Text frames list representative genes for the specific modules. (C) Bar plot showing the percentage of gene pairs within each module that display species-specific expression correlation. Bars are colored by functional categories (see the listed pathways). Two modules enriched for the “Carbon metabolism” pathway serve as the reference. Significance was assessed using Fisher’s exact test (* FDR < 0.05; ** FDR < 0.0001).

### 2.5 Convergent transcriptional regulation in human and mouse spermatogenesis

While LS lncRNAs cause cross-species regulatory differences, they also allow many developmental processes to converge on conserved outcomes. Since mammalian spermatogenesis is a precise developmental process but how the process is regulated by LS lncRNAs remains unclear, we investigated the regulation of spermatogenesis by LS lncRNAs using single-cell RNA-seq (scRNA-seq) datasets from Sertoli (ST), spermatogonia (SG), spermatocytes (SC), round spermatids (RS), and elongated spermatids (ES) cells in humans and mice (Murat et al., 2023). Across these cell types, we identified 599 PS/SS/HS lncRNAs and 1,196 RS/MS lncRNAs. Several LS lncRNAs exhibit distinct tissue-specific expression in humans, including *LINC01356* in the testis and brain, and *WASIR2* in the ovary and brain, suggesting their species-specific global impacts. To focus on the regulation of the transcriptional core of spermatogenesis, we identified 395 conserved spermatogenesis marker genes, which are the intersection of (a) highly expressed genes (HEG) within at least one cell type, (b) highly variable genes (HVG) across the five cell types, and (c) one-to-one orthologous PCGs in human, chimpanzee, mouse, opossum, and platypus. These markers define a deeply conserved transcriptional program, and many of them, including *ZPBP* (whose protein binds the oocyte zona pellucida) and *MYO6* (which regulates actin reorganization during spermatogenesis from *C. elegans* to mammals), were not targeted in previous studies (Murat et al., 2023; Shami et al., 2020).

To examine how 599 PS/SS/HS lncRNAs and 1,196 RS/MS lncRNAs regulate these conserved markers, we predicted their DBSs in promoter regions of the 395 markers (Supplementary Table 9). Then, we used the *eGRAM* program to identify transcriptional regulatory modules in each cell type based on both DBS targeting and expression correlation relationships between lncRNAs and their targets. Each module comprises a regulator set (LS lncRNAs) and a target set (conserved markers). Initially, 8, 18, 13, 4, 2, and 10, 8, 14, 12, 17 modules were identified in ST, SG, SC, RS, and ES cells in humans and mice, reflecting species-specific regulation. After merging modules until no two shared≥98% of target genes, each cell type in both species converged on a single dominant module with essentially the same markers (Fig. 5A; Supplementary Figs. 12-14). Thus, despite substantial differences in initial modules, the final modules in humans and mice converge on nearly identical core genes. To evaluate whether final modules faithfully represent cell identity, we built five independent reference marker sets derived from humans and opossums (Murat et al., 2023), and compared the ten marker sets in the final modules in humans and mice with the five reference sets. Eight marker sets showed the highest similarity (Jaccard index) to their corresponding reference sets, and two showed the second-highest similarity, supporting the faithfulness of the inferred modules.

**Figure 5.**
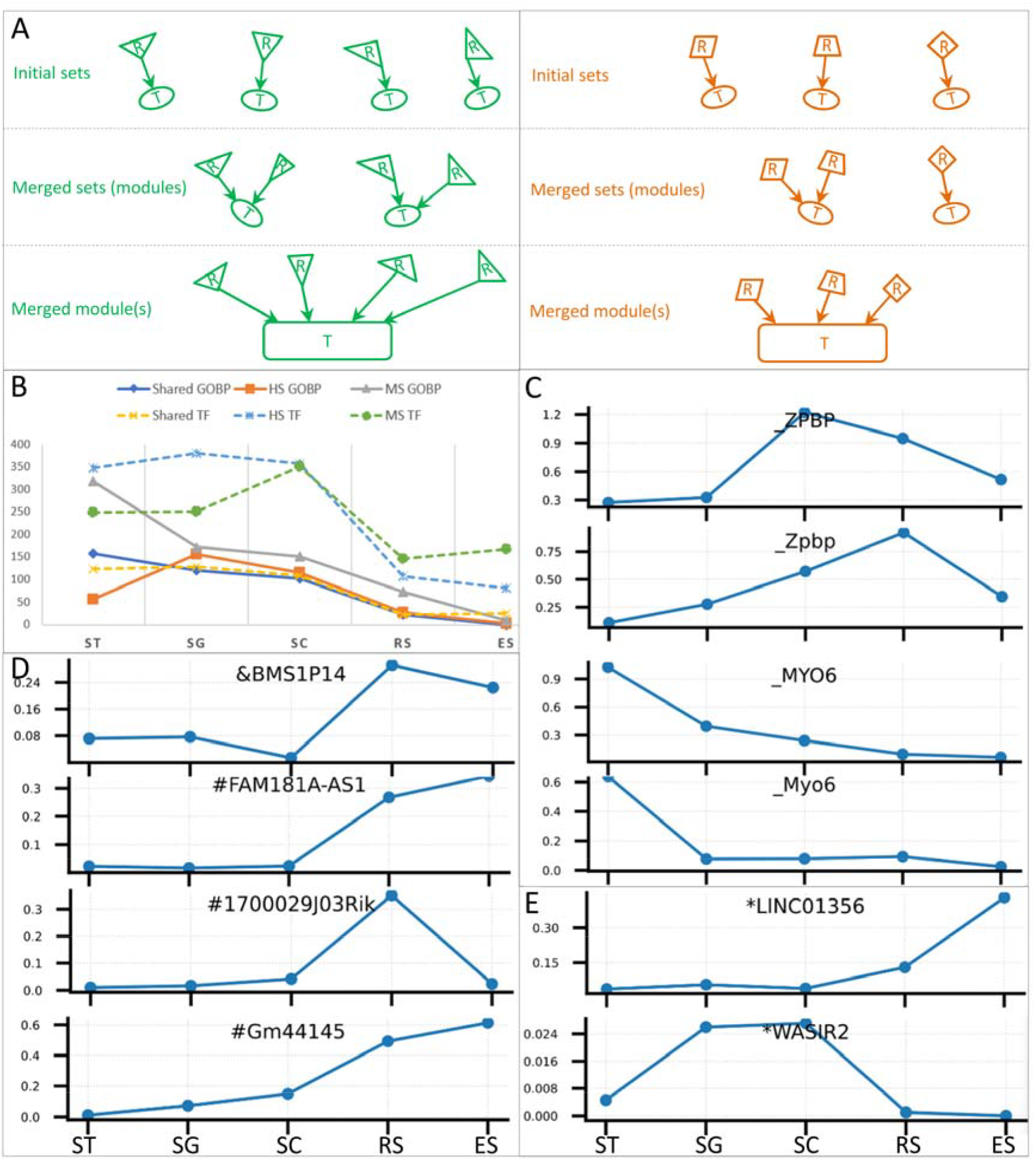
Transcriptional regulation by LS lncRNAs in human and mouse spermatogenesis. (A) Illustration of initial sets, merged sets, and the final module in each cell type in mice and humans. ‘R’ and ‘T’ denote the regulator and target sets, respectively. (B) Numbers of shared, human-specific, and mouse-specific GO and TRANSFAC terms identified in merged sets in the five cell types. (C) Expression dynamics of two conserved markers in humans and mice. (DE) Expression dynamics of several LS lncRNAs.

We then performed ORA for markers in each module to identify shared, human-specific, and mouse-specific GO terms and TRANSFAC terms. Of note, many terms detected in early cell types (ST and SG) were species-specific, and the number of terms decreased progressively from ST to ES cells (Fig. 5B; Supplementary Fig. 12). This gradual decrease mirrors the increasing developmental commitment during spermatogenesis, as transcriptional programs become progressively constrained toward terminal cell identity. Expression dynamics of LS lncRNAs and marker genes reveal coordinated yet distinctly regulated programs. Marker genes exhibit expected stage-specific patterns (Fig. 5C), whereas LS lncRNAs are distinctly up- or down-regulated in late stages (Fig. 5DE), indicating the integration of LS lncRNAs into species- and stage-specific programs. The proportion of markers that show correlated expression with LS lncRNAs, and the number of LS lncRNAs that are expressed, increase as spermatogenesis progresses.

Finally, we assessed whether enriched GO terms reflect established mechanisms of spermatogenesis. The majority of enriched terms reasonably correspond to canonical processes such as meiosis, chromatin remodeling, and germ cell differentiation. In addition, GO terms that have drawn limited attention in the mainstream spermatogenesis literature are supported by related findings, including “circadian rhythm” (GO:0007623) that was reported to regulate spermatogonial differentiation (Liu et al., 2025b), “microautophagy” (GO:0016237) that participates in chromatoid body homeostasis in haploid round spermatids (Wang et al., 2022), and “cellular response to thyroglobulin and triiodothyronine” (GO:1904017) that reflects the involvement of triiodothyronine (T3) in germ cell differentiation and survival (Morenas et al., 2024). Collectively, the spermatogenesis analysis illustrates how LS lncRNAs facilitate developmental processes that converge on conserved outcomes while generating cross-species regulatory differences.

## 3. Discussion

### 3.1 Systematic Mapping of Reference Non-Coding Genomes

LncRNAs regulate gene expression, immune responses, and drug sensitivity (Clapes et al., 2021; Guo et al., 2021; Zhao et al., 2021), and substantial lncRNAs are lineage-specific. However, previous studies examined lineage specificity using *de novo* transcript assembly from organ-derived RNA-seq data across multiple species and using sequence-only alignment tools to infer noncoding orthology. Nor did they examine related TEs, DBDs, and DBSs. It remains unclear which GENCODE-annotated lncRNAs are lineage-specific and why, whether they cause cross-species differences in transcriptional regulation, and how they facilitate developmental processes that converge on conserved outcomes. This study systematically identifies the evolutionary origins of GENCODE-annotated lncRNAs in humans and mice, their DBDs and DBSs, and the embedded TEs. This approach represents a significant conceptual and methodological departure from prior studies. We demonstrate that LS TEs critically determine LS lncRNAs and their DBDs and DBSs. This structural link causes extensive, lineage-specific rewiring of transcriptional networks without disrupting deeply conserved, core cellular machineries. The majority of HS lncRNA DBSs are located within human-mouse syntenic blocks, suggesting the rewiring primarily emerges through the structural modification and “tinkering” of pre-existing genomic architecture rather than through the *de novo* creation of new genomic sequences.

### 3.2 Bridging the Preclinical Translational Gap

Our results provide a long-awaited mechanistic explanation for translational failures in preclinical animal models (Drake, 2013; Mak et al., 2014; Seok et al., 2013). When target genes exhibit highly species-specific distributions across signaling pathways and tissue transcriptomes, a therapeutic target validated in mice may have no functional equivalent in humans. Our analyses of multiple high-profile datasets illustrate this non-coding mismatch. Taking *LIN28B* as an example. In human cancers, the widespread, TE-driven ectopic expression of the oncogenic lncRNA *LIN28B* is guided by simian-specific TEs embedded within its promoter region (Jang et al., 2019). In mice, however, this promoter region contains rodent-specific TEs. This suggests that the ectopic activation of this oncogene follows species-specific regulatory logic that cannot be faithfully modeled in rodents. Taking *EPIC1* as another example. Exons of the lncRNA *EPIC1*—which mediates tumor immune evasion and immunotherapy resistance (Guo et al., 2021)—overlap multiple simian-specific TEs. This suggests that *EPIC1* is subject to human-specific regulatory constraints and helps explain why therapeutic responses targeting this axis in mice may not translate to human clinical trials. Our results provide a framework to map GENCODE lncRNAs directly to clinical databases and separate preclinical disease models into functionally convergent versus divergent archetypes.

### 3.3 Evolutionary Robustness and Path Dependency

Our results also help reveal how regulatory divergence generates developmental conservation. In spermatogenesis, while humans and mice employ entirely distinct repertoires of LS lncRNAs, these divergent regulators converge on nearly identical, deeply conserved core marker genes to drive equivalent cell-fate determination. This explains a profound evolutionary duality: the rapid genomic turnover of TE-derived lncRNAs provides the extraordinary regulatory flexibility required for lineages to adapt and innovate, while the ultimate targeting of conserved protein-coding core networks ensures developmental robustness (Senft and Macfarlan, 2021; Thybert et al., 2018).

The structural reliance of LS lncRNAs on LS TEs fundamentally dictates the trajectory of evolutionary innovation. Because specific TE families accumulate in massive, non-random bursts within distinct lineages, the subsequent birth of LS lncRNAs and their DBSs is intrinsically, structurally biased. In this sense, the TE landscape of a specific species constrains the available architectural blueprints for future regulatory networks and preordains certain evolutionary novelties. For example, the rapid accumulation of young TEs in primates provides a pre-existing sequence reservoir that systematically directs the emergence of primate-specific regulatory networks (Batzer and Deininger, 2002; Osmanski et al., 2023). The PS lncRNAs *LINC01356* and *WASIR2*, which are expressed exclusively in the human brain and reproductive tissues, demonstrate how lineage-specific expansion of specific TEs can preordain the emergence of highly complex, multi-trait innovations.

### 3.4 Limitations and Future Directions

First, as genome annotation remains in progress, LS lncRNAs may be underestimated. For example, our pipeline did not identify *LINC01876*—a recently reported, LINE1-derived, HS lncRNA expressed exclusively during brain development (Garza et al., 2023). Second, the identified LS lncRNAs also await downstream analysis, such as the human/mouse comparative pan-cancer analysis (Lin et al., 2026a). Third, because ncRNA-DNA triplexes follow specific base-pairing rules, DBS-like sequences naturally occur across different genomes. This implies that LS lncRNAs have the physical capacity to bind to non-LS DBSs under artificial conditions, such as in foreign knock-in experiments (Liu et al., 2025a).

## 4. Materials and Methods

### 4.1 Identification of LS lncRNAs

The *Infernal* program performs RNA homology searches using covariance models that integrate primary sequence and secondary structure information (Nawrocki and Eddy, 2013; Nawrocki et al., 2009). We previously conducted exon-level searches for 13,562 GENCODE-annotated human lncRNA genes (GENCODE V18) across 16 representative mammalian genomes using *Infernal* (Lin et al., 2019; Lin et al., 2026b). To improve orthology inference, exon searches were constrained within syntenic regions determined by UCSC pairwise genome alignments, and the aligned regions were extended at both ends. If more than 50% of the exons of a human lncRNA gene had detectable matches in a target genome, the gene was considered to have an ortholog in that genome. Since full-length lncRNA transcripts often exhibit rapid sequence evolution and variable exon structures, and no universally accepted threshold exists for lncRNA orthology inference, the majority-exon criterion provides a conservative and reproducible operational definition. Using the same strategy, we searched for orthologs of 13,450 GENCODE-annotated mouse lncRNA genes (GENCODE M22) in four mammalian genomes: *Mus caroli* (CAROLI_EIJ_v1.1), *Mus pahari* (PAHARI_EIJ_v1), rat (*Rnor_6*.*0*), and rabbit (*OryCun2*.*0*). These rodent genomes represent key divergence nodes within Muridae, while the rabbit was included as a non-rodent Glires outgroup to distinguish rodent-restricted from more broadly conserved Glires lncRNAs.

A human lncRNA gene was defined as human-specific (HS) if no ortholog was detected in any of the 16 surveyed mammals. A mouse lncRNA gene was defined as mouse-specific (MS) if no ortholog was detected in any of the four surveyed mammals. The selected primate and rodent genomes provide comparable phylogenetic resolution, as divergence between primates (e.g., human–orangutan) and rodents (e.g., mouse–rat) reflects similar evolutionary timescales (Thybert et al., 2018). At the time of the study, high-quality genome assemblies were available for more primate species than rodent species; however, these chosen rodent genomes span major divergence nodes within Rodentia, providing sufficient resolution for lineage-specific classification. Using genome builds GRCh37 and GRCm38, we identified 66 HS lncRNA genes and 212 MS lncRNA genes. Coordinates of HS lncRNAs were converted from GRCh37 to GRCh38 using *liftOver* (Raney et al., 2024).

To further resolve lineage specificity in primates, we examined the presence of orthologs of human lncRNA exons in three simians (chimpanzee, macaque, marmoset), three prosimians (tarsier, mouse lemur, tree shrew), and the mouse. The mouse was used as a phylogenetically appropriate outgroup for primate-specific classification because rodents are more closely related to primates than to other mammalian clades such as Laurasiatheria. A human lncRNA was defined as primate-specific (PS) if it had orthologs in at least one simian and one prosimian but not in mouse, as simian-specific (SS) if it had orthologs in at least one simian but not in any prosimian and mouse, and as human-specific (HS) if it had no orthologs in any other species. A mouse lncRNA was defined as rodent-specific (RS) if it had orthologs in at least one rodent species (*Mus caroli, Mus pahari*, or rat) but not in rabbit and human, and mouse-specific (MS) if it had no orthologs in any other species. In the spermatogenesis datasets, lineage- and species-specific lncRNAs were annotated with distinct symbols: ‘&’ for PS, ‘#’ for SS and RS, and ‘*’ for HS and MS.

### 4.2 Prediction of lncRNA DBDs and DBSs

LncRNA-DNA binding follows experimentally characterized base-pairing rules to form RNA-DNA triplexes (Abu Almakarem et al., 2012), thereby allowing lncRNAs to recruit epigenetic modification enzymes to their binding sites (Chu et al., 2011; Engreitz et al., 2016; Yap et al., 2010). Multiple methods predict lncRNA triplex-targeting sites (TTSs) using substring search based on these rules (Buske et al., 2012; Kuo et al., 2019), but this approach has three limitations: (a) TTSs are very short and disconnected, whose reliability needs additional statistical tests, (b) TTSs are poor at capturing densely populated lncRNAs at binding sites, and (c) triplex-forming oligonucleotides (TFOs) are not accurately identified. *LongTarget* was designed to overcome these limitations by simultaneously predicting DBDs and DBSs. It translates the DNA sequence into 24 RNA sequences using 24 integrated rule sets, applies a modified Smith–Waterman local alignment algorithm to 24 pairs of RNA sequences to identify all high-scoring local alignments, identifies a set of densely overlapping triplexes (TFOs and TTSs) that define paired DBD and DBS, and ranks DBDs (He et al., 2015; Lin et al., 2019). The alignment can accommodate flexible distributions of mismatches, insertions, and deletions in triplexes and allows *binding affinity = length × identity* (*the percentage of matched base pairs*) to be used to compare DBSs within and across lncRNAs. The Fasim-*LongTarget* version solves the drawback of high computational time (Wen et al., 2022).

For each HS and MS lncRNA, DBSs were predicted within 5-kb promoter regions (defined as 3500 bp upstream to 1500 bp downstream of the transcription start site, TSS) of 179,128 Ensembl-annotated human transcripts (GRCh38.79) and 95,052 Ensembl-annotated mouse transcripts (GRCm38.87) using *LongTarget* (*Length* = 50, *Identity* = 60%) (Fig. 1A–D). TSS annotations were obtained directly from Ensembl. For each PS, SS, and RS lncRNA, DBSs were predicted within the 10-kb promoter region of conserved spermatogenesis marker genes using *Fasim*-*LongTarget* (the same parameters) (Murat et al., 2023). For each marker gene, we selected the most representative TSS from the UCSC EPDnew Promoters track, which integrates TSS annotations from HGNC, GENCODE, Ensembl, RefSeq, FANTOM, UCSC, and ENCODE (Dreos et al., 2017). The two programs typically predict multiple DBDs per lncRNA, each associated with a set of DBSs, and rank DBDs according to their cumulative DBS support. Since DBD1—the top-ranked DBD—consistently exhibits substantially greater DBS coverage than other DBDs, which suggests stronger statistical support, our downstream analyses focused on DBD1s and their DBSs to ensure robustness and comparability across lncRNAs. Importantly, the biological relevance of DBD1s has been experimentally validated. Using CRISPR/Cas9, we deleted DBD1 in multiple lncRNAs across several cell lines, performed RNA sequencing before and after DBD1 deletion, analyzed differentially expressed genes (DEGs), and conducted functional assays demonstrating altered cellular phenotypes (He et al., 2026; Lin et al., 2026b). Benchmarking based on real data also supports the reliability and biological relevance of the predicted DBD1–DBS interactions (Lin et al., 2026b; Liu et al., 2017; Wen et al., 2022).

### 4.3 Determination of TEs’ origins and contributions to genes and regulatory elements

#### 4.3.1 Determination of TE Origins

To infer the evolutionary origins of TEs embedded in HS and MS lncRNAs, we developed a computational phylogenomic pipeline to identify homologous TE sequences across 87 representative species. These species span major evolutionary clades, including primates, rodents, Lagomorpha, Laurasiatheria, other placental mammals, marsupials, monotremes, birds and reptiles, fishes, chordates, arthropods, nematodes, and mollusks, thereby enabling inference of the phylogenetic distribution and minimum evolutionary age of each TE. The pipeline integrates: (a) three BLAST-based search tools: NCBI BLAST API, NCBI SRA Toolkit (blastn_vdb), and local BLAST (Altschul et al., 1990), (b) multiple sequence databases: NCBI Nucleotide collection (nt), Whole-Genome Shotgun contigs (WGS), Transcriptome Shotgun Assembly (TSA), and reference genomes of human (GRCh38) and mouse (GRCm38), (c) *MAFFT* for multiple sequence alignment (Katoh and Standley, 2013), (d) *RAxML* for phylogenetic reconstruction (Stamatakis, 2014).

For each TE sequence, the following steps were performed. (a) A BLASTn search was submitted through the NCBI BLAST API against the Nucleotide collection (nt) database (*reward* = 2, *penalty* = −3, *gap opening cost* = 5, *gap extension cost* = 2, and *E-value threshold* = 0.05). (b) A standalone BLASTn search was performed using blastn_vdb (SRA Toolkit) against WGS and TSA datasets of the selected 87 species. (c) Local BLASTn searches were conducted against the human (GRCh38) and mouse (GRCm38) reference genomes to ensure complete detection of paralogous or fragmented copies. (d) To standardize similarity scores across TEs of varying lengths, raw BLAST bit-scores were normalized by dividing by the maximal theoretical score of the query TE. Under the scoring scheme, the maximal score equals 2 × TE length. (e) Hits with normalized scores greater than 0.4 were retained as qualified homologs, while lower-scoring hits were discarded. This threshold filters out weak or fragmented matches while retaining moderately diverged homologs. The E-value threshold was set to a permissive value to maximize sensitivity at the initial retrieval stage, and specificity was enforced by the normalized score cutoff. Empirical inspection of known lineage-restricted elements (e.g., simian SINEs) showed that the 0.4 threshold retains expected lineage patterns. (f) The UCSC REST API was used to determine whether qualified hits overlapped annotated gene exons in the respective genomes.

After homolog collection, multiple sequence alignments were generated using *MAFFT* (default parameters). For phylogenetic reconstruction, we retained (a) the top five homologs in human and mouse and (b) the top one homolog per other species. Since humans and mice were the focal genomes and we aimed to evaluate TE contributions to both lncRNAs and PCGs within these species, retaining multiple homologs allows assessment of representative TE copies in related genes while avoiding excessive redundancy from highly expanded TE families. In contrast, for non-focal species, only presence or absence was required for evolutionary inference; therefore, a single representative homolog per species was sufficient. Alignments were trimmed using *BMGE* (default parameters) to remove poorly aligned or highly divergent regions (Criscuolo and Gribaldo, 2010). Maximum-likelihood phylogenetic trees were then constructed using *RAxML* (substitution model = ‘GTRGAMMA’, 1000 bootstrap replicates, mechanism = ‘raxmlHPC-HYBRID-SSE3’, threads = (TE length / 500) + 2). Bootstrap support was interpreted cautiously for short elements, and trees were visualized using *ggtree*. TE origin was inferred from the most basal clade in which qualified homologs were detected, as supported by phylogenetic clustering. For example, TEs with homologs restricted to simians were classified as simian-specific (SS), whereas those present across rodents but absent outside Rodentia were classified as rodent-specific (RS). We emphasize that “origin” refers to the earliest detectable phylogenetic distribution under the defined similarity threshold and does not exclude the possibility of highly diverged ancestral copies.

#### 4.3.2 Determination of TE contributions to genes and regulatory elements

Annotated TE coordinates were obtained from RepeatMasker annotations for the GRCh38 and GRCm38 genomes. To estimate TE contributions to LS lncRNAs and their regulatory elements, we intersected TE coordinates with: lncRNA exons, DBD1 regions, and DBSs. If a TE overlapped an exon, DBD, or DBS by >30% of its length, the TE was considered to have contributed to that element.

To assess TE contributions to PCGs, exon coordinates were obtained from Ensembl Release 101 annotations (GRCh38 and GRCm38), comprising 391,676 human exons and 256,983 mouse exons. Each exon sequence was subjected to the same phylogenomic pipeline described above to identify homologs across the 87 species. Exons with qualified homologs (normalized score >0.4) exclusively in simians but absent in all other species were classified as simian-specific (SS) exons. Similarly, exons with qualified homologs restricted to rodents were classified as rodent-specific (RS) exons. If an SS or RS exon overlapped a TE of corresponding lineage by >30% of its length, the exon was considered to have received contribution from that TE. This operational definition refers to TE-involved exonization events, in which lineage-restricted TEs contribute sequence to lineage-specific coding exons.

### 4.4 Analysis of preferential binding and lineage-specific rewiring

#### 4.4.1 Preferential binding between LS lncRNAs and LS DBSs

Unlike inferring the evolutionary origin of TEs in lncRNAs by searching for TE sequences across 87 species, we determined the lineage of TEs in DBSs by performing homologous genomic mapping of DBS sequences using *liftOver* (Lin et al., 2014; Pehrsson et al., 2019; Raney et al., 2024). This is to examine whether lncRNAs containing TEs of a given lineage preferentially bind DBSs containing TEs of the same lineage rather than DBSs containing TEs of other lineages. Since spermatogenesis is a conserved process intensively regulated by lncRNAs, it provides a stringent context for analyzing the preferential binding. First, we predicted DBSs of 1,795 LS lncRNAs in 395 conserved spermatogenesis marker genes in humans and mice (Section 4.2). To determine lineages of TEs in human DBSs, TE coordinates in the GRCh38 genome were converted to the chimpanzee (panTro6), mouse lemur (micMur2), mouse (GRCm38), and rat (rn7) genomes using *liftOver* (*minMatch* = 0.1, as in previous studies (Pehrsson et al., 2019)). A TE was classified as: (a) simian-specific if it had counterparts in chimpanzee but not in mouse lemur, (b) primate-specific if it had counterparts in both chimpanzee and mouse lemur but not in mouse or rat, (c) conserved in mammals if it had counterparts in mouse and/or rat. Similarly, TE coordinates in the GRCm38 genome were converted to the rat, chimpanzee, and human genomes. A TE was classified as: (a) rodent-specific if it had counterparts in rat but not in human or chimpanzee, (b) conserved in mammals if it had counterparts in human and/or chimpanzee. Failure to obtain a *liftOver* mapping indicates a lack of detectable homology.

After the TE classification, an LS TE was considered to contribute to an LS DBS if it overlapped by >30% of the DBS length. We then tested whether lineage-matched lncRNA–DBS pairs (i.e., both containing TEs of the same lineage) were more frequent and/or exhibited stronger predicted binding affinity than lineage-mismatched pairs. Differences in binding strength were assessed using the two-sided Mann–Whitney U test, and differences in counts were evaluated using the Chi-square test.

#### 4.4.2 Lineage-specific rewiring of transcriptional regulation

To further evaluate lineage-specific rewiring at the genomic level, we examined whether human DBSs fall within human–mouse syntenic blocks. Syntenic block information was obtained from the NCBI Comparative Genome Viewer (CGV) using the GRCh38–GRCm38 genome alignment. CGV classifies genomic regions into: (a) SP (Syntenic Position): regions located within conserved syntenic blocks, (b) NoHit: regions without detectable aligned counterparts, (c) OffChrm: regions aligning to a different chromosome, (d) Mix: regions partially overlapping syntenic and non-syntenic segments. NoHit, OffChrm, or Mix regions reflect disruptions or rearrangements in positional conservation. A DBS was considered to reside within a syntenic block if it fell within SP regions. *liftOver* was used to determine how many human DBSs have mouse counterparts (*minMatch* = 0.9, as DBSs are short functional elements). Together, synteny classification and high-stringency *liftOver* mapping were used to assess the extent of lineage-specific regulatory rewiring.

### 4.5. Functional enrichment analysis

#### 4.5.1 GO term analysis

GO enrichment analysis was performed using *g:Profiler* and *PANTHER* to assess functional biases among target genes of HS and MS lncRNAs (Mi et al., 2019; Raudvere et al., 2019). To evaluate the functional impacts of HS and MS lncRNAs genome-wide, target genes were ranked in descending order according to the number of regulatory lncRNAs. To focus on genes most strongly influenced by LS lncRNAs and reduce background noise from weakly targeted genes, the top 10% of ranked genes were selected for enrichment analysis. Over-representation analysis (ORA) was conducted using *g:Profiler* with the “*ordered query*” option enabled, which incorporates the ranking information of the input gene list. GO BP terms with an FDR < 0.05 were considered significant (Fig. 2A).

To directly compare functional biases between HS and MS lncRNA regulatory targets, ORA was also performed using *PANTHER* (FDR < 0.05). In this analysis, the top 10% target genes of HS lncRNAs and the top 10% target genes of MS lncRNAs were contrasted to identify GO BP terms that are relatively over-represented in one species compared with the other. This cross-species comparison was designed to highlight functional divergence between human- and mouse-specific lncRNA regulatory networks rather than to assess enrichment against a whole-genome background. Enriched and under-represented GO BP terms were identified accordingly (Fig. 2B).

To further examine functional divergence associated with TE age, ORA was performed on target genes of HS lncRNAs in humans whose DBDs were derived from different Alu subfamilies (AluY-, AluS-, and AluJ-derived). Enrichment of GO BP terms (FDR < 0.05) was assessed using the same analytical framework (Fig. 2C).

#### 4.5.2. KEGG pathway analysis

We estimated the species specificity of molecular signaling potentially regulated by HS and MS lncRNAs by analyzing 382 KEGG pathways from the KEGG database (https://www.genome.jp/kegg). We examined whether genes containing HS or MS lncRNA DBSs, as well as genes whose exons overlap simian- or rodent-specific TE sequences, show lineage-biased distributions across pathways. For each gene within a pathway, we calculated the proportions of HS and MS lncRNAs containing DBSs in that gene; for each pathway, these proportions were summed across all member genes to generate a pathway-level regulatory score (Fig. 2D). Pathways with higher cumulative scores indicate greater predicted regulatory input from HS and MS lncRNAs. Similarly, for each pathway, we calculated the proportions of genes whose exons overlap simian- or rodent-specific TE sequences, generating a pathway-level estimate of lineage-specific TE contribution. These analyses provide a comparative assessment of pathway-level regulatory and TE-associated biases between humans and mice.

### 4.6. Combined analysis of gene expression profiles and signaling pathways

To investigate transcriptomic impacts of HS and MS lncRNAs and LS TEs, we performed integrative analyses across tissues and species.

#### 4.6.1 Correlation between HS and MS lncRNAs and target genes

We first examined whether transcriptional regulation predicted by DBS binding is associated with cross-species and cross-tissue variation in gene expression. Gene expression matrices and *bigWig* files for 54 human GTEx tissues and 17 mouse tissues were obtained from recount3 (Consortium et al., 2017; Schaum et al., 2020). Gene-level counts were converted to TPM values and log-transformed prior to analysis. For each tissue, Spearman correlation coefficients between HS and MS lncRNAs and their predicted target genes were computed using the *spearmanr* function in the *scipy* package. A lncRNA–gene pair was considered significantly correlated in a tissue if |ρ| > 0.3 and FDR < 0.05 (Fig. 3AB). Predicted DBS binding and expression correlation were integrated to provide complementary evidence consistent with putative regulatory relationships between HS and MS lncRNAs and their targets. For each tissue, we calculated the proportion of HS and MS lncRNAs that exhibited significant expression correlation with at least one predicted target gene. Differences in these proportions across human and mouse tissues were used to estimate tissue-specific divergence in lineage-associated transcriptional regulation (Fig. 3AB).

#### 4.6.2 Expression analysis of TE-derived exons

To assess the transcriptomic contribution of LS TE-derived exons within HS and MS lncRNA target genes, we extracted exon-level read coverage from *bigWig* files using the *pyBigWig* package (https://github.com/deeptools/pyBigWig). Read counts were converted to TPM values and log-transformed. For each tissue, the median TPM across all samples was used to represent exon expression. To evaluate tissue biases, median TPM values for each exon were standardized within species to z-scores across tissues. An exon with a z-score > 1 was considered relatively highly expressed in that tissue.

#### 4.6.3 Cross-species comparison of one-to-one orthologous genes

To quantify cross-species transcriptional divergence, one-to-one orthologous genes between human and mouse in the same 11 tissues were obtained from the Ensembl Compara API (Vilella et al., 2009). For each tissue, gene expression was represented by the median TPM across samples. When a tissue contained subregions, samples from all subregions were combined. Within each species, median TPM values were standardized to z-scores across tissues to allow comparison of relative expression patterns. For each gene and tissue, cross-species expression divergence was defined as: |Δz|=|z_human_−z_mouse_|. This metric captures species-specific deviations in tissue-relative expression patterns.

#### 4.6.4 Extension to pathway-level divergence

To evaluate pathway-level and tissue-level combined transcriptional divergence, the |Δz| value was computed for each gene in each KEGG pathway and tissue. This enabled comparison of cross-species expression differences across pathways and tissues. To assess pathway- and tissue-specific divergence, the distribution of |Δz| values for genes within a given pathway and tissue was compared with the distribution of |Δz| values for genes across all pathways in the pancreas using a one-sided Mann–Whitney U test (testing whether |Δz|_specific pathway+specific tissue_ > |Δz|_all pathway+pancreas_). P-values were adjusted for multiple testing (FDR < 0.001) (Fig. 3C). The pancreas was selected as an empirical baseline because it exhibited the lowest proportion of HS and MS lncRNA–target correlation events and minimal enrichment of LS TE-associated regulation, indicating comparatively weak lineage-associated transcriptomic divergence.

### 4.7 Analysis of transcriptional regulation by HS and MS lncRNAs in AD

To examine whether LS lncRNA regulatory networks are associated with cross-species differences in AD, we applied *eGRAM* to RNA-seq data from human AD patients and mouse AD models. RNA-seq data from the lateral temporal lobe of AD patients (n = 12) and from the hippocampus of late-onset AD mouse models (n = 14) were obtained from published studies (Baglietto-Vargas et al., 2021; Nativio et al., 2018). Although the sampled brain regions are not anatomically identical, both regions are strongly implicated in AD pathology and neurodegeneration. The input gene set comprised genes (median TPM > 0.1) in AD-related KEGG pathways. The human and mouse input matrices contained matched one-to-one orthologous genes, enabling cross-species comparison. Regulatory modules were identified based on three thresholds: DBS binding affinity ≥60, Spearman correlation ≥0.3, and module size ≥5 genes.

After module identification (Fig. 4), pairwise expression correlations among target genes within each module were computed using the *spearmanr* function in the *scipy* package. P-values were adjusted for multiple testing using the FDR method. A correlation between two target genes in a human module was defined as human-specific if: (a) the correlation in human AD samples satisfied |ρ| > 0.5 and FDR < 0.05, and (b) the correlation between the corresponding mouse orthologs in mouse AD samples satisfied |ρ| < 0.1 and FDR > 0.05. Conversely, a correlation between two target genes in a mouse module was defined as mouse-specific if: (a) the correlation in mouse AD samples satisfied |ρ| > 0.5 and FDR < 0.05, and (b) the correlation between the corresponding human orthologs satisfied |ρ| < 0.1 and FDR > 0.05. Thus, species-specific correlations were defined as strong and statistically significant in one species but weak and non-significant in the other. Because previous analyses indicated that metabolism-related pathways exhibit minimal DBS enrichment and weak association with HS and MS lncRNAs (Fig. 4AB), gene pairs within these pathways were used as an empirical low-regulation reference set. For both human and mouse, the proportion of species-specific correlated gene pairs within metabolism-related modules was calculated as background. For each AD-related module, the proportion of species-specific correlated gene pairs was compared with the corresponding background proportion using Fisher’s exact test. Statistical significance was assessed to determine whether nervous system– and immune-related AD modules exhibited elevated species-specific co-expression relative to the low-regulation reference (Fig. 4C).

### 4.8 Development of the *eGRAM* program for analyzing transcriptional regulation in single cells

Most existing approaches for transcriptional regulatory analysis cluster genes into disjoint modules based on correlated expression. Such approaches do not explicitly model the many-to-many relationships between regulators and targets and the physical evidence of regulatory targeting. The original *GRAM* (Gene Regulatory Analysis Model) was designed to overcome these limitations by identifying overlapping regulatory relationships and exploring both correlated expression and TF-TFBS targeting (Bar-Joseph et al., 2003). However, it does not incorporate lncRNA regulation and cannot analyze scRNA-seq data. Our developed *eGRAM* retains key features of *GRAM* while enabling integration of lncRNA–DNA binding information. *eGRAM* for bulk RNA-seq data requires these inputs: (a) an RNA-seq expression matrix, (b) an lncRNA–DBS binding matrix, and (c) (optional) a TF–DBS binding matrix (He et al., 2026). Key parameters include: (a) lncRNA binding affinity threshold, (b) TF binding affinity threshold, (c) expression correlation threshold, and (d) minimum module size. The expression correlation between regulators and targets is computed using Pearson or Spearman correlations; regulatory modules are constructed by integrating binding evidence and expression correlations; and genes may participate in multiple modules.

scRNA-seq data present additional challenges, including high sparsity (dropout) and nonlinear gene–gene relationships. Therefore, the *eGRAM* version (v3) for scRNA-seq data employs a different strategy for estimating correlations. *eGRAMv3* operates on log-normalized counts without data imputation. Instead of Pearson or Spearman correlation, it employs the Maximal Information Coefficient (MIC) from the MINE (Maximal Information-based Nonparametric Exploration) family of statistics (Reshef et al., 2011). The MIC algorithm measures the maximum normalized mutual information achievable across all possible grid discretizations of the paired variables and can detect both linear and nonlinear relationships. MIC returns two statistics: (a) MIC (maximal information coefficient), (b) TIC (total information coefficient). Because MIC values depend on sample size and grid resolution, large MIC values alone do not guarantee statistical significance. The original MINE framework uses permutation testing to assess significance; however, permutation testing is computationally impractical for scRNA-seq datasets with thousands of genes and cells.

The majority of gene pairs in high-dimensional transcriptomic datasets are expected to lack statistically detectable association, reflecting the sparsity of biological regulatory networks and co-expression structure observed in both bulk and single-cell studies (Barabasi and Oltvai, 2004; Basso et al., 2005). Empirical analyses of gene co-expression networks consistently show that only a small fraction of all possible gene pairs exhibit meaningful association, while most pairs display near-zero correlation (Andrews and Hemberg, 2018; Skinnider et al., 2019; Svensson, 2020). Under this sparsity assumption, MIC values can be viewed as arising from a mixture of a dominant null component (background) and a small non-null component (correlated). We therefore revised the MIC algorithm by employing adaptive thresholding using empirical null-distribution modeling. Our revised algorithm extracts the upper-triangular values from the MIC matrix, fits a two-component Gaussian mixture model, uses the “mean + 3 standard deviation” or “mean + 4 standard deviation” of the background as a threshold, and identifies significantly correlated gene pairs based on the chosen threshold. Simulation results indicate that this algorithm tolerates dropouts and reliably detects associations when the sample size is sufficiently large (hundreds of cells), enabling scalable significance estimation without computationally intensive permutations.

To evaluate performance, we compared MIC-based correlation detection with multiple alternative methods, including Hoeffding’s D, maximal correlation, distance correlation, Bayesian correlation, and BigSur (Hoeffding, 1948; Nguyen et al., 2014; Sanchez-Taltavull et al., 2020; Silkwood et al., 2024), using both simulated and real scRNA-seq datasets. Simulated datasets consisted of 10 genes and 100 cells, with predefined correlated gene pairs of varying strengths and varying sparsity levels (zero proportions ranging from 70% to 90%). Across simulations: when zero proportion <80%, MIC detected both directly specified and indirectly correlated gene pairs with high sensitivity; when zero proportion reached 90%, detection power decreased substantially for all methods; in the 80–90% sparsity range, combining MIC and TIC improved detection, recovering over 60% of predefined correlated gene pairs. Across tested scenarios, the revised MIC algorithm consistently showed greater sensitivity for nonlinear and sparse associations than alternative metrics, particularly under moderate-to-high dropout rates.

To improve computational efficiency for large scRNA-seq datasets, we implemented two optimizations. First, we used an adaptive *alpha* to control the *for*-loop that finds the grid that maximizes the mutual information between variables. In the MIC algorithm, *alpha* is a hyperparameter that defines the maximum number of grid partitions the algorithm searches through to compute the MIC value, relative to the sample size. The maximal grid size is bounded by *Partition*(n) = n^*alpha*^, where *n* is the sample size and *alpha* is commonly set to 0.6. We empirically evaluated *alpha* values between 0.45 and 0.6 and found that they produced stable results for medium-to-large scRNA-seq datasets while substantially reducing computation time. Second, we parallelized the MIC algorithm. For a human spermatogenesis dataset (thousands of cells), the parallel implementation completed within approximately 20 hours on a single AMD 8845HS CPU with 64 GB RAM.

### 4.9. Analysis of human and mouse spermatogenesis

#### 4.9.1 Obtaining spermatogenesis scRNA-seq data

We downloaded scRNA-seq datasets covering spermatogenesis across multiple mammalian species from the EBI BioStudies repository (https://www.ebi.ac.uk/biostudies/arrayexpress/studies/) (Murat et al., 2023). In the original datasets, outliers and potential doublets had been removed, gene expression was normalized and log-transformed, and gene annotations from Ensembl Release 87 were used. We extracted genes with Ensembl gene symbols, including 28,727 in humans (GRCh38.87), 15,351 in chimpanzees (CHIMP2.1.4.87), and 21,911 in mice (GRCm38.87). For each species, we retained Sertoli (ST), spermatogonia (SG), spermatocytes (SC), round spermatids (RS), and elongated spermatids (ES) as defined by the original cell-type annotations. Cells with poor quality were further filtered using the interquartile range (IQR) criterion (lower bound = Q1 − 1.5 IQR; upper bound = Q3 + 1.5 IQR). For each gene and each cell type, we computed the mean expression level. Highly expressed genes (HEGs) were defined based on mean expression within each cell type. Genes were then ranked by the range of mean expression between the maximum and minimum across the five cell types, and the most highly variable genes (HVGs) were selected.

To identify conserved and LS genes involved in spermatogenesis, orthologous gene relationships among humans, chimpanzees, mice, opossums, and platypuses were retrieved from Ensembl BioMart (Release 112). Among 191,508 annotated genes across the five species, 8,565 exhibited one-to-one orthology. Conserved spermatogenesis markers were defined as the intersection of: (a) the top 200 HEGs in each cell type, (b) HVGs, and (c) one-to-one orthologs across the five species (Supplementary Table 10). LS genes included both protein-coding and lncRNA genes. LS protein-coding genes were identified following Kirilenko et al. (https://genome.senckenberg.de//download/TOGA/) (Kirilenko et al., 2023), which classified orthologous genes across placental mammals and birds into categories such as “many2zero,” “one2one,” and “one2zero” using pairwise genome comparisons (e.g., GRCh38–GRCm38, GRCm38– GRCh38). In GRCh38–GRCm38 comparisons, “many2zero” and “one2zero” genes correspond to PS/SS and HS genes relative to the mouse. In GRCm38–GRCh38 comparisons, “many2zero” and “one2zero” genes correspond to RS and MS genes relative to the human.

LS lncRNAs were identified by searching for orthologs of GENCODE-annotated human and mouse lncRNAs across multiple mammalian genomes (Section 4.1; Supplementary Tables 8, 9). LS lncRNAs with consistent gene symbols between Ensembl Releases 87 and 101 and expressed during spermatogenesis were retained (Supplementary Table 10). Conserved and lineage-specific (PS, SS, HS, RS, MS) lncRNAs were labeled using specific symbols ( ‘_’ indicating conserved in mammals, ‘&’ indicating PS, ‘#’ indicating SS and RS, and ‘*’ indicating HS and MS) in the input files, enabling evaluation of transcriptional regulation by LS lncRNAs. Previous studies identified spermatogenesis markers using Seurat’s *FindAllMarkers* function after dividing pseudotime trajectories into 20 bins (Murat et al., 2023; Shami et al., 2020). However, many such markers (e.g., *TEX11*) are not conserved across mammals, limiting cross-lineage comparison between primates and rodents. Under our definition, primate-specific spermatogenesis markers such as *TEX11* and *TEX41* were not classified as conserved markers.

#### 4.9.2 Analyzing transcriptional regulation by LS lncRNAs in spermatogenesis

Using *eGRAMv3* and the above genes and datasets, we analyzed transcriptional regulation mediated by PS/SS/HS lncRNAs in humans and RS/MS lncRNAs in mice to examine how lineage-specific regulatory mechanisms may produce conserved spermatogenic outcomes.

First, we adopted the following parameter settings. (a) *zero_proportion* = 0.85. Genes with zero expression in >85% of cells were filtered out. A lower threshold substantially reduced gene coverage, whereas thresholds ≥90% impaired MIC performance under high sparsity. (b) The *binding affinity* threshold = 36. Because regulation was determined jointly by DBS evidence and expression correlation, relatively permissive thresholds were selected to avoid over-constraining inference. Also, the binding affinity of multiple RS/MS lncRNAs in mouse *Zpbp* is 35-40. (c) The threshold for correlated expression in the MIC algorithm. The threshold was defined as the “mean + 3 standard deviations” of the background component. Correlations can be considered significant under either the setting of “mic-or-tic” (either MIC or TIC significant) or the setting of “mic-and-tic” (both MIC and TIC significant). At the gene level, the two settings yielded 5,708 and 4,898 significant correlations (i.e., TRUEs), respectively. At the “merged set” level (Fig. 5A), the two settings returned largely the same results (significant correlations), “mic-or-tic” generated slightly more GO terms, including multiple cohesion-related GO terms, than “mic-and-tic”, yet the top GO terms and all KEGG pathways were the same. Since Cohesin loading onto chromatin is a crucial process in spermatogenesis, we used the “mic-or-tic” setting. (d) *Module size* = 50. Only regulatory modules containing ≥50 target genes were reported. (e) FDR threshold = 0.01 for pathway enrichment analysis. (f) The *alpha* parameter for MIC computation. The *alpha* was 0.6 if the sample size < 500, 0.55 if < 1000, 0.5 if < 1500, and 0.45 if ≥ 1500. We observed that the fixed *alpha* (0.6) and the adaptive *alpha* (with the above values) yielded essentially the same results.

Second, we validated modules. To assess whether the identified final modules reasonably indicate cell types, we constructed five reference marker sets (ST, SG, SC, RS, ES) based on the intersections of marker genes in humans and opossums (Murat et al., 2023). For each final module, we computed the Jaccard index between its target gene set and each reference marker set. If the target gene set and the corresponding reference marker set have the highest Jaccard Index, then it indicates that the final module reasonably matches the cell type.

Third, the identification of transcriptional regulatory modules followed these steps. (a) For each LS lncRNA, identify its collaborator lncRNAs based on expression correlation, and this generates a “regulator set”. (b) For each regulator set, compute correlations between lncRNAs therein and all other genes and evaluate DBS binding. Genes satisfying both DBS binding and correlation criteria were defined as the “target set” (“initial sets” in Fig. 5A). The regulator set and its corresponding target set constituted a module. (c) Merge two regulator sets if they share identical target sets (“merged sets” in Fig. 5A). (d) Determine whether target genes (conserved spermatogenesis marker genes) in each module are significantly enriched for genes in GO terms, TRANSFAC terms, and KEGG/Wikipathway pathways. (e) Iteratively merge modules whose target gene sets overlapped by >98% of target genes. In each round, if more than two modules have target sets that share the same percentage of genes, the two modules whose lncRNA sets share the highest percentage of lncRNAs are merged (“merged modules” in Fig. 5A). (f) Compute the Jaccard index between the target set in each final module and the five reference marker sets.

## Acknowledgements

This work was supported by the National Natural Science Foundation of China (31771456) and the China Postdoctoral Science Foundation (2020M682788).

## Author contributions

H.Z. designed the study; P.P. proposed the TE analysis pipeline. H.Z. and J.L. developed the *eGRAM* version. J.L. and Y.W. performed most of the analyses; H.Z. performed the spermatogenesis analysis; J.Z., W.X., and S.H. contributed to the experimental validation of the lncRNA-DBS prediction. H.Z. wrote the manuscript. All authors have read the manuscript and consent to its publication.

## Additional information

Supplementary file 1: Supplementary figures 1-14.

Supplementary file 2: Supplementary tables 1-14.

Related manuscript: Lin et al. Lineage-specific lncRNAs critically determine cross-species differences in tumors. bioRxiv 2026 (doi: https://doi.org/10.64898/2026.05.01.722350).

