## Supplementary figures for "Intrinsically linked lineage specificity of transposable elements, lncRNA genes, and transcriptional regulation" (from another version of this preprint: version 2, 2026-05-10)

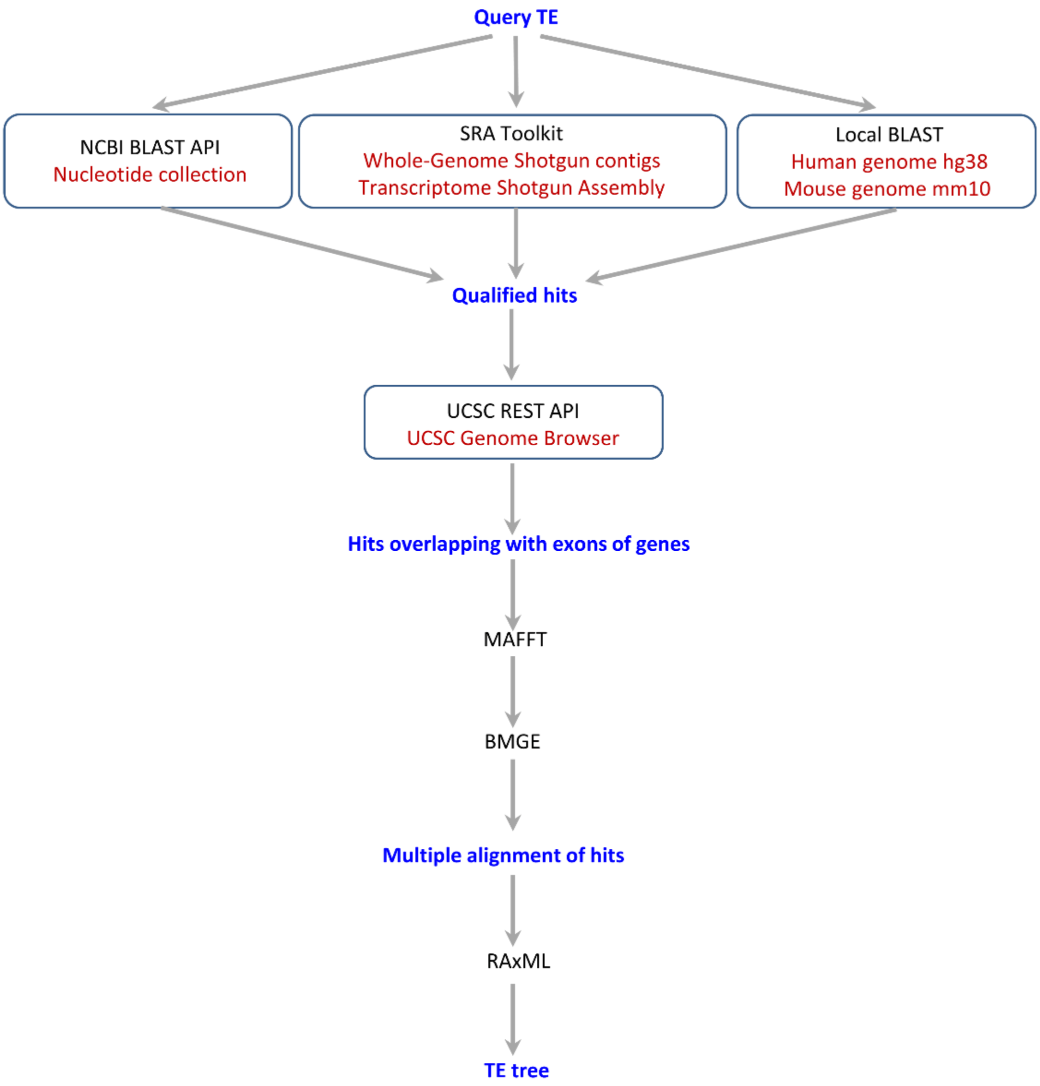


Supplementary Figure 1. The computational pipeline for searching homologues of TEs in the 87 species and building phylogenetic trees of the hits. This pipeline was also used to search for homologues of protein-coding genes' exons in the 87 species. Blue items indicate inputs, outputs, and results; black items indicate computational programs; red items indicate data resources.

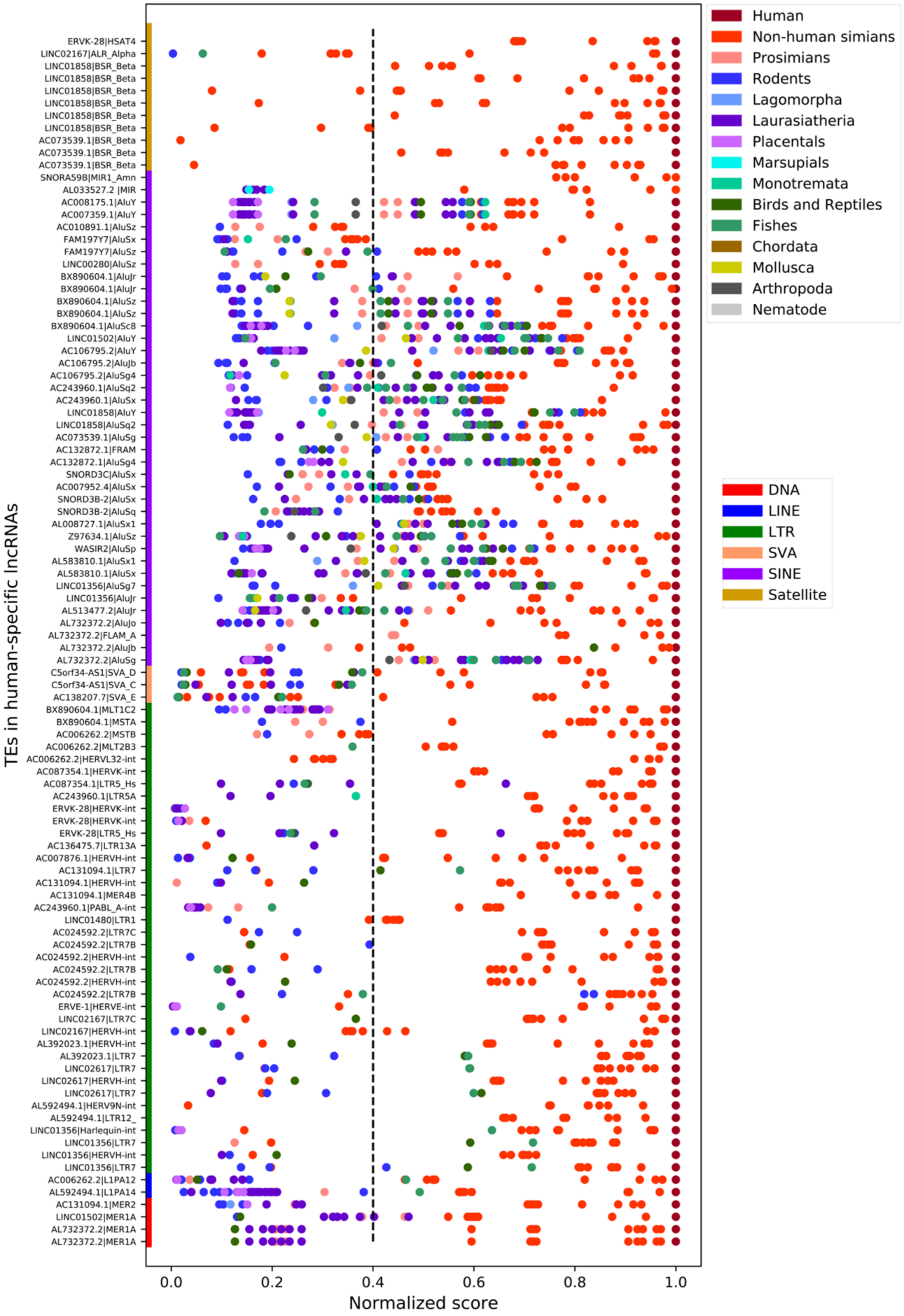
ERVK-28|HSAT4
LINC02167|ALR_Alpha
LINC01858|BSR_Beta
LINC01858|BSR_Beta
LINC01858|BSR_Beta
LINC01858|BSR_Beta
LINC01858|BSR_Beta
LINC01858|BSR_Beta
AC073539.1|BSR_Beta
AC073539.1|BSR_Beta
AC073539.1|BSR_Beta
SNORA59B|MIR1_Amn
AL033527.2 |MIR
AC008175.1|AluY
AC007359.1|AluY
AC010891.1|AluSz
FAM197Y7|AluSx
FAM197Y7|AluSz
LINC00280|AluSz
BX890604.1|AluJr
BX890604.1|AluJr
BX890604.1|AluSz
BX890604.1|AluSz
BX890604.1|AluSc8
LINC01502|AluY
AC106795.2|AluY
AC106795.2|AluJb
AC106795.2|AluSg4
AC243960.1|AluSq2
AC243960.1|AluSx
LINC01858|AluY
LINC01858|AluSq2
AC073539.1|AluSg
AC132872.1|FRAM
AC132872.1|AluSg4
SNORD3C|AluSx
AC007952.4|AluSx
SNORD3B-2|AluSx
SNORD3B-2|AluSq
AL008727.1|AluSx1
Z97634.1|AluSz
WASIR2|AluSp
AL583810.1|AluSx1
AL583810.1|AluSx
LINC01356|AluSg7
LINC01356|AluJr
AL513477.2|AluJr
AL732372.2|AluJo
AL732372.2|FLAM_A
AL732372.2|AluJb
AL732372.2|AluSg
C5orf34-AS1|SVA_D
C5orf34-AS1|SVA_C
AC138207.7|SVA_E
BX890604.1|MLT1C2
BX890604.1|MSTA
AC006262.2|MSTB
AC006262.2|MLT2B3
AC006262.2|HERVL32-int
AC087354.1|HERVK-int
AC087354.1|LTR5_Hs
AC243960.1|LTR5A
ERVK-28|HERVK-int
ERVK-28|HERVK-int
ERVK-28|LTR5_Hs
AC136475.7|LTR13A
AC007876.1|HERVH-int
AC131094.1|LTR7
AC131094.1|HERVH-int
AC131094.1|MER4B
AC243960.1|PABL_A-int
LINC01480|LTR1
AC024592.2|LTR7C
AC024592.2|LTR7B
AC024592.2|HERVH-int
AC024592.2|LTR7B
AC024592.2|HERVH-int
AC024592.2|LTR7B
ERVE-1|HERVE-int
LINC02167|LTR7C
LINC02167|HERVH-int
AL392023.1|HERVH-int
AL392023.1|LTR7
LINC02617|LTR7
LINC02617|HERVH-int
LINC02617|LTR7
AL592494.1|HERV9N-int
AL592494.1|LTR12_
LINC01356|Harlequin-int
LINC01356|LTR7
LINC01356|HERVH-int
LINC01356|LTR7
AC006262.2|L1PA12
AL592494.1|L1PA14
AC131094.1|MER2
LINC01502|MER1A
AL732372.2|MER1A
AL732372.2|MER1A
TEs in human-specific lncRNAs
0.0
0.2
0.4
0.6
0.8
1.0
Normalized score
Human
Non-human simians
Prosimians
Rodents
Lagomorpha
Laurasiatheria
Placentals
Marsupials
Monotremata
Birds and Reptiles
Fishes
Chordata
Mollusca
Arthropoda
Nematode
DNA
LINE
LTR
SVA
SINE
Satellite

Supplementary Figure 2. The distribution of hits of query TEs (TEs in HS lncRNAs) in the 87 species that cover 15 classes and clades. Hit scores are normalized, and the dashed vertical line indicates the threshold for identifying qualified hits. Only hits reaching 50% of the length of Dfam-annotated consensus sequences of the corresponding TE subfamilies are shown. Most hits (especially those of TEs of the SINE subfamily) have scores >0.4 only in simians.

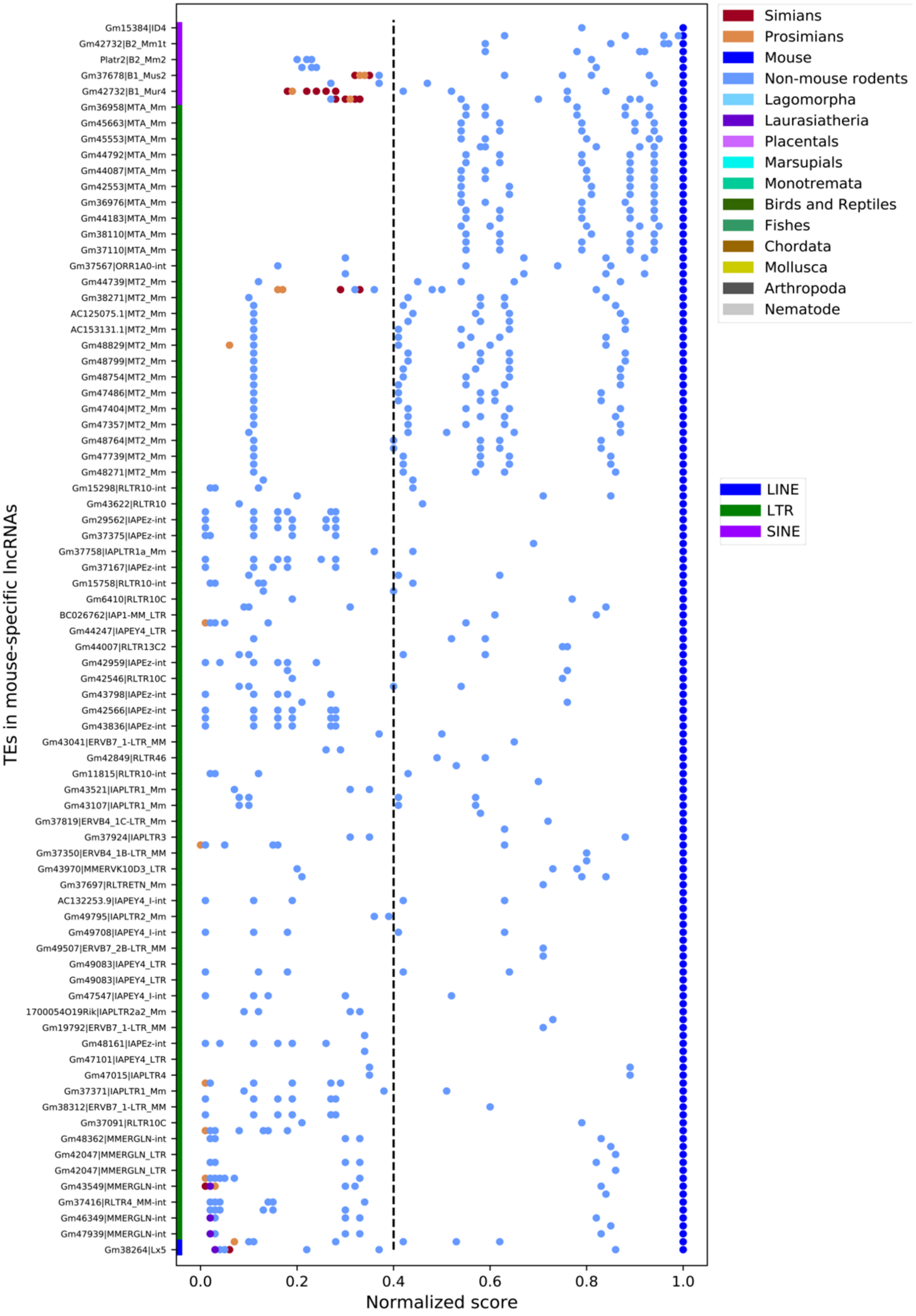
TEs in mouse-specific lncRNAs
Gm15384|ID4
Gm42732|B2_Mm1t
Platr2|B2_Mm2
Gm37678|B1_Mus2
Gm42732|B1_Mur4
Gm36958|MTA_Mm
Gm45663|MTA_Mm
Gm45553|MTA_Mm
Gm44792|MTA_Mm
Gm44087|MTA_Mm
Gm42553|MTA_Mm
Gm36976|MTA_Mm
Gm44183|MTA_Mm
Gm38110|MTA_Mm
Gm37110|MTA_Mm
Gm37567|ORR1A0-int
Gm44739|MT2_Mm
Gm38271|MT2_Mm
AC125075.1|MT2_Mm
AC153131.1|MT2_Mm
Gm48829|MT2_Mm
Gm48799|MT2_Mm
Gm48754|MT2_Mm
Gm47486|MT2_Mm
Gm47404|MT2_Mm
Gm47357|MT2_Mm
Gm48764|MT2_Mm
Gm47739|MT2_Mm
Gm48271|MT2_Mm
Gm15298|RLTR10-int
Gm43622|RLTR10
Gm29562|IAPEz-int
Gm37375|IAPEz-int
Gm37758|IAPLTR1a_Mm
Gm37167|IAPEz-int
Gm15758|RLTR10-int
Gm6410|RLTR10C
BC026762|IAP1-MM_LTR
Gm44247|IAPEY4_LTR
Gm44007|RLTR13C2
Gm42959|IAPEz-int
Gm42546|RLTR10C
Gm43798|IAPEz-int
Gm42566|IAPEz-int
Gm43836|IAPEz-int
Gm43041|ERVB7_1-LTR_MM
Gm42849|RLTR46
Gm11815|RLTR10-int
Gm43521|IAPLTR1_Mm
Gm43107|IAPLTR1_Mm
Gm37819|ERVB4_1C-LTR_Mm
Gm37924|IAPLTR3
Gm37350|ERVB4_1B-LTR_MM
Gm43970|MMERVK10D3_LTR
Gm37697|RLTRETN_Mm
AC132253.9|IAPEY4_I-int
Gm49795|IAPLTR2_Mm
Gm49708|IAPEY4_I-int
Gm49507|ERVB7_2B-LTR_MM
Gm49083|IAPEY4_LTR
Gm49083|IAPEY4_LTR
Gm47547|IAPEY4_I-int
1700054O19Rik|IAPLTR2a2_Mm
Gm19792|ERVB7_1-LTR_MM
Gm48161|IAPEz-int
Gm47101|IAPEY4_LTR
Gm47015|IAPLTR4
Gm37371|IAPLTR1_Mm
Gm38312|ERVB7_1-LTR_MM
Gm37091|RLTR10C
Gm48362|MMERGLN-int
Gm42047|MMERGLN_LTR
Gm42047|MMERGLN_LTR
Gm43549|MMERGLN-int
Gm37416|RLTR4_MM-int
Gm46349|MMERGLN-int
Gm47939|MMERGLN-int
Gm38264|Lx5
0.0
0.2
0.4
0.6
0.8
1.0
Normalized score
Simians
Prosimians
Mouse
Non-mouse rodents
Lagomorpha
Laurasiatheria
Placentals
Marsupials
Monotremata
Birds and Reptiles
Fishes
Chordata
Mollusca
Arthropoda
Nematode
LINE
LTR
SINE

Supplementary Figure 3. The distribution of hits of query TEs (TEs in MS lncRNAs) in the 87 species that cover 15 classes and clades. Hit scores are normalized, and the dashed vertical line indicates the threshold for identifying qualified hits. Only hits reaching 50% of the length of Dfam-annotated consensus sequences of the corresponding TE subfamilies are shown. Nearly all hits are in rodents.

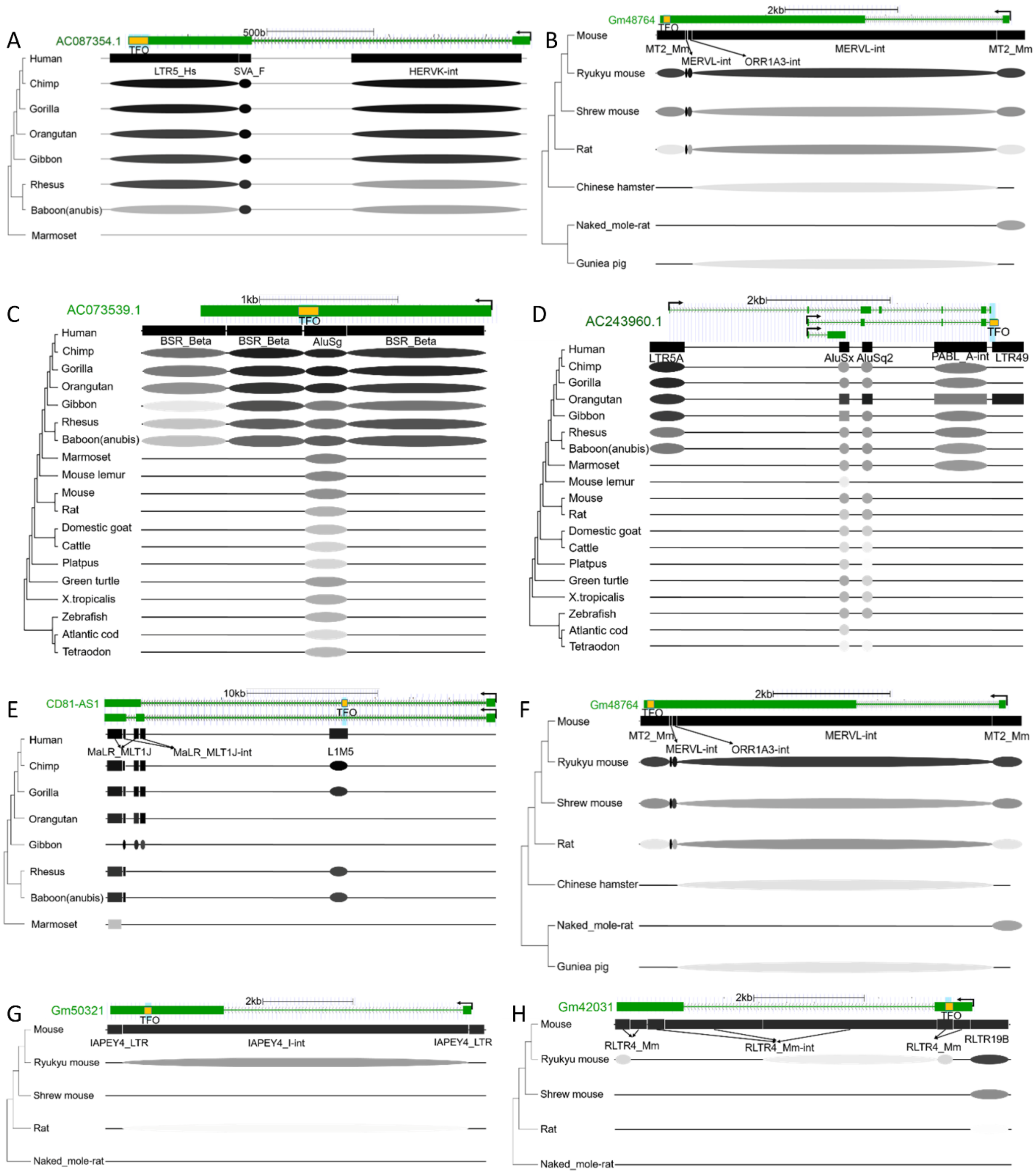


Supplementary Figure 4. Phylogenetic trees of query TEs and their hits. Shown are some trees that reveal the evolutionary origins of TEs in HS/MS lncRNAs. Rectangles and ellipses indicate hits at orthologous and non-orthologous genomic positions. Greyscales indicate normalized scores. (A) The tree of TEs in AC087354.1 and their hits, indicating that these TEs are simian-specific. (B) The tree of TEs in Gm48764 and their hits, suggesting that these TEs are rodent-specific. (C) The tree of TEs in AC073539.1 and their hits, suggesting that one TE is conserved but others are simian-specific. (D) The tree of TEs in AC243960.1 and their hits, suggesting that two TEs are conserved but others are simian-specific. (E) The tree of TEs in CD81-AS1 and their hits, suggesting that these TEs are simian-specific. (F) The tree of TEs in Gm48764 and their hits. (G) The tree of TEs in Gm50321 and their hits. (H) The tree of TEs in Gm42031 and their hits.

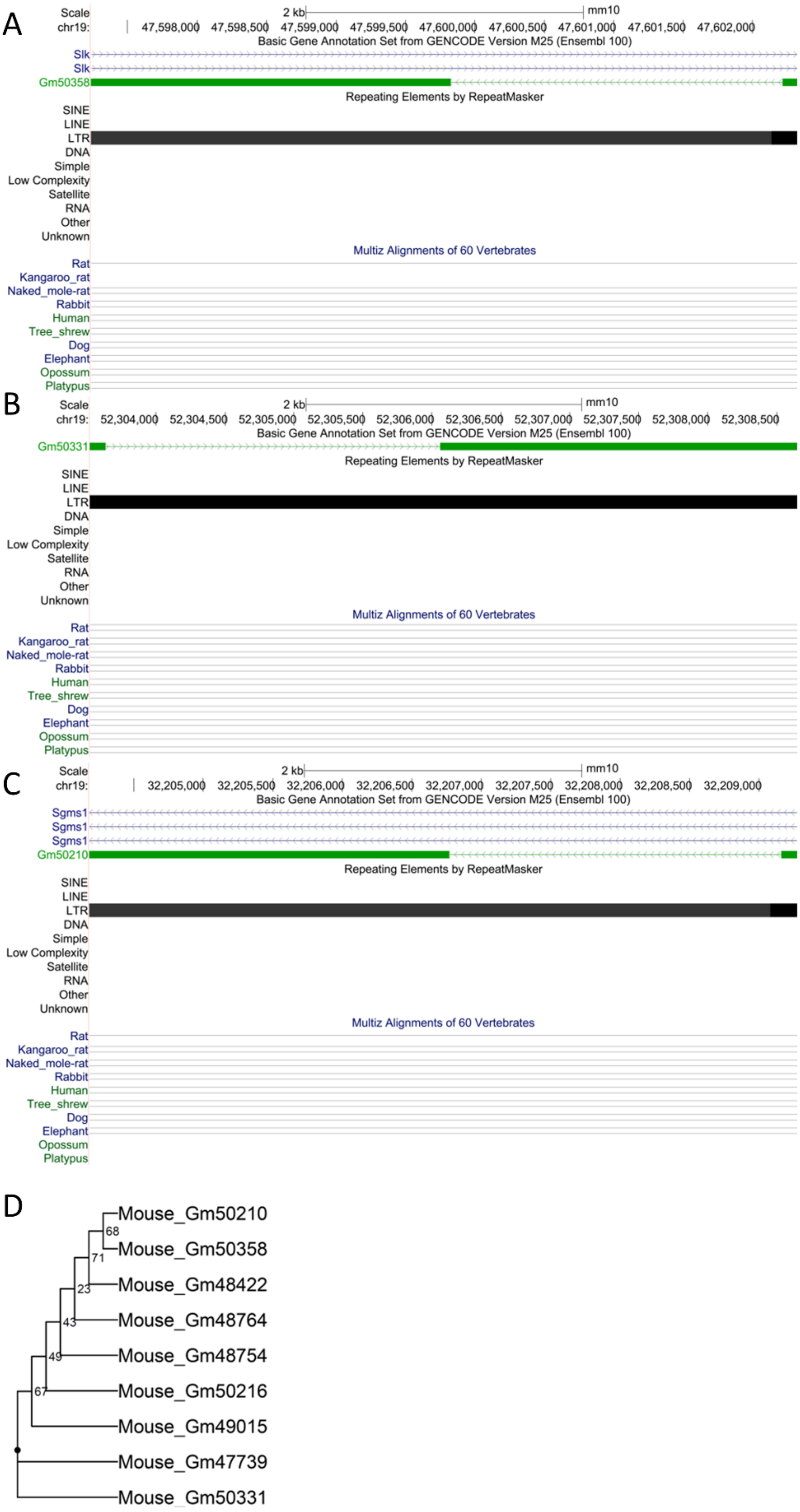
A
Scale
chr19:
2 kb
mm10
47,598,000 47,598,500 47,599,000 47,599,500 47,600,000 47,600,500 47,601,000 47,601,500 47,602,000
Basic Gene Annotation Set from GENCODE Version M25 (Ensembl 100)
Slk
Slk
Gm50358
Repeating Elements by RepeatMasker
SINE
LINE
LTR
DNA
Simple
Low Complexity
Satellite
RNA
Other
Unknown
Multiz Alignments of 60 Vertebrates
Rat
Kangaroo_rat
Naked_mole-rat
Rabbit
Human
Tree_shrew
Dog
Elephant
Opossum
Platypus
B
Scale
chr19:
2 kb
mm10
52,304,000 52,304,500 52,305,000 52,305,500 52,306,000 52,306,500 52,307,000 52,307,500 52,308,000 52,308,500
Basic Gene Annotation Set from GENCODE Version M25 (Ensembl 100)
Gm50331
Repeating Elements by RepeatMasker
SINE
LINE
LTR
DNA
Simple
Low Complexity
Satellite
RNA
Other
Unknown
Multiz Alignments of 60 Vertebrates
Rat
Kangaroo_rat
Naked_mole-rat
Rabbit
Human
Tree_shrew
Dog
Elephant
Opossum
Platypus
C
Scale
chr19:
2 kb
mm10
32,205,000 32,205,500 32,206,000 32,206,500 32,207,000 32,207,500 32,208,000 32,208,500 32,209,000
Basic Gene Annotation Set from GENCODE Version M25 (Ensembl 100)
Sgms1
Sgms1
Sgms1
Gm50210
Repeating Elements by RepeatMasker
SINE
LINE
LTR
DNA
Simple
Low Complexity
Satellite
RNA
Other
Unknown
Multiz Alignments of 60 Vertebrates
Rat
Kangaroo_rat
Naked_mole-rat
Rabbit
Human
Tree_shrew
Dog
Elephant
Opossum
Platypus
D
Mouse_Gm50210
68
Mouse_Gm50358
71
Mouse_Gm48422
23
Mouse_Gm48764
43
Mouse_Gm48754
49
Mouse_Gm50216
67
Mouse_Gm49015
Mouse_Gm47739
Mouse_Gm50331

Supplementary Figure 5. LTR/ERVL in an MS lncRNA contributes to 9 paralogous MS lncRNAs. (A-C) The genomic view of 3 of the 9 paralogous MS lncRNA genes (the genomic views of the remaining 6 paralogues are similar). (D) The phylogenetic tree built using the hits of the LTR/ERVL in Gm50331. These hits are detected only in mice, suggesting that the LTR/ERVL is mouse-specific.

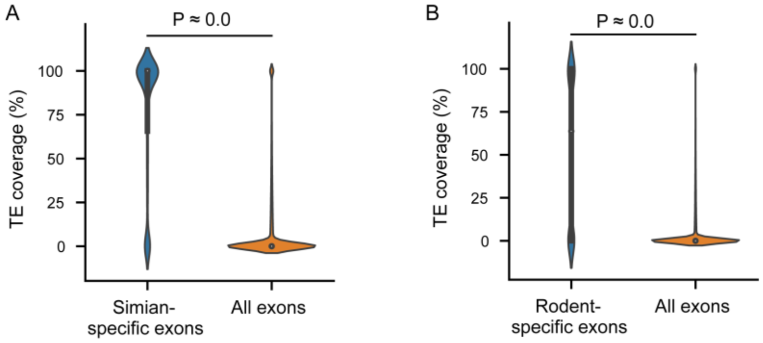


Supplementary Figure 6. The TE coverage of lineage-specific exons compared with the TE coverage of all Ensembl-annotated exons. Each exon's TE coverage is calculated by dividing the TE-overlapping length by the total length. Compared with all Ensembl-annotated exons of protein-coding genes in human and mouse genomes, these simian/rodent-specific exons have significantly higher TE coverage ($P \approx 0$ for both simian-specific and rodent-specific exons, one-sided Mann–Whitney test). (A) The TE coverage of simian-specific exons. (B) The TE coverage of rodent-specific exons.

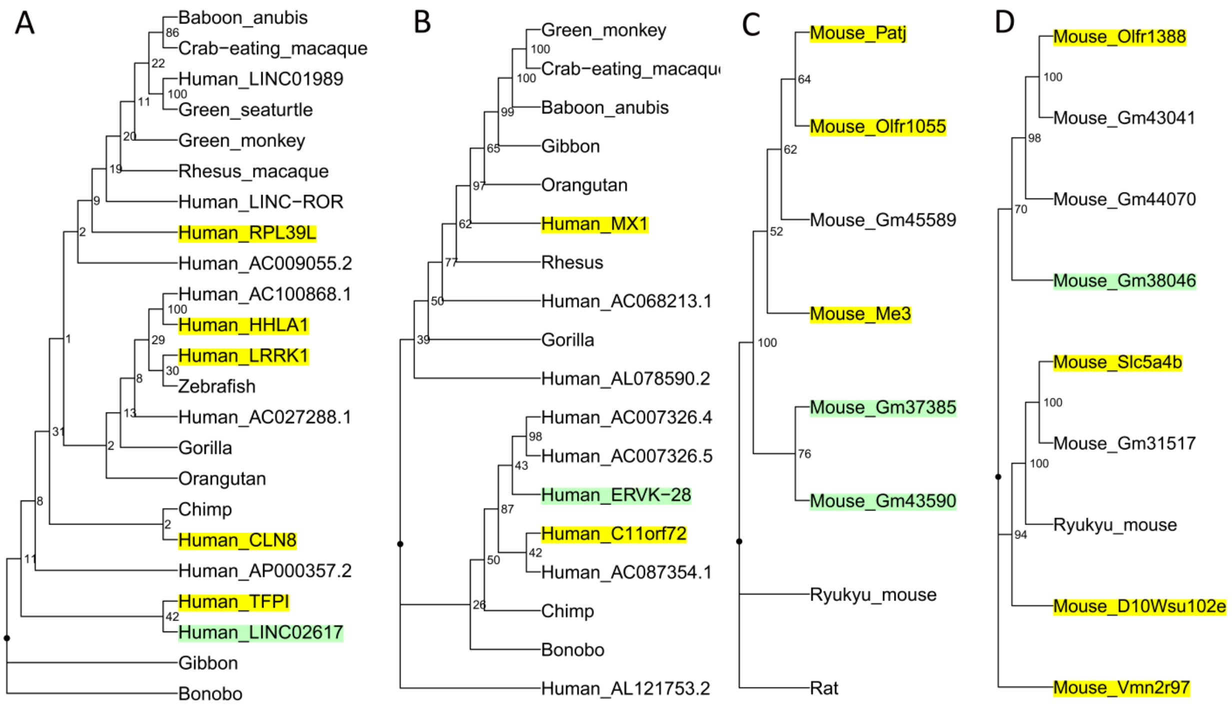


Supplementary Figure 7. TEs that contribute sequences to exons of HS/MS lncRNA genes also contribute sequences to exons of protein-coding genes. In humans, many of these protein-coding genes are essential for the neural and immune systems (e.g., *LRRK1* and *CLN8* are associated with Parkinson's disease and neural development, and *MX1* participates in the cellular antiviral response). In mice, some of these protein-coding genes encode mouse-specific olfactory receptors (e.g., *Olfr1055* and *Olfr1388*). In these phylogenetic trees, query TEs are colored in green, and their hits that overlap with exons of protein-coding genes by >20 bp are colored in yellow. Note that nodes such as "Human_LRRK1" indicate the hit of a TE in the human gene *LRRK1*, but nodes such as "Green_monkey" and "Gibbon" indicate hits in the Green_monkey and Gibbon genomes. (A) The phylogenetic tree of LTR/EVR1 in LINC02617 and its hits. (B) The phylogenetic tree of LTR/ERVK in ERVK-28 and its hits. (C) The phylogenetic tree of LTR/ERVK in Gm37385 and Gm43590 and its hits. (D) The phylogenetic tree of LTR/ERVK in Gm38046 and its hits.

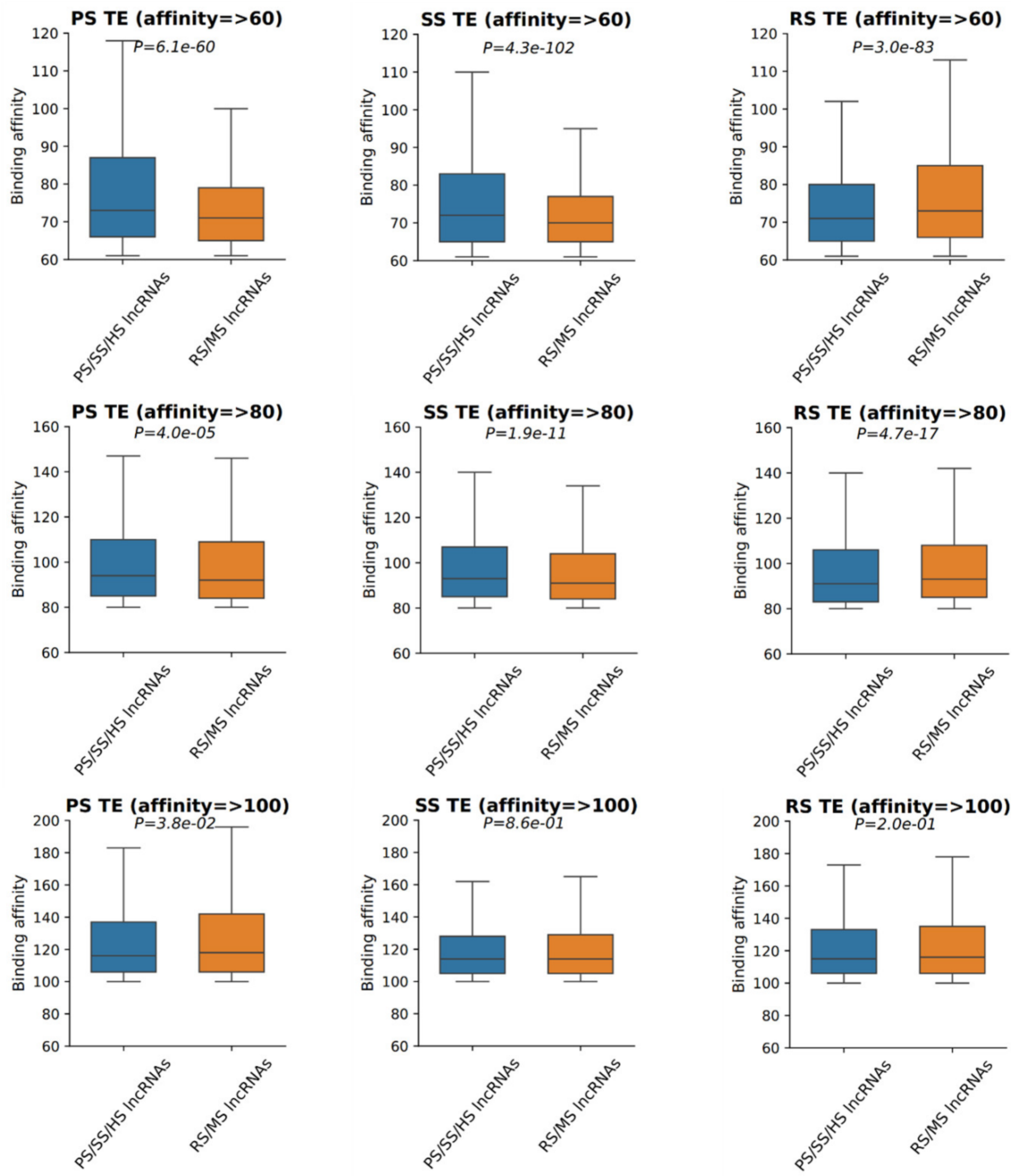


Supplementary Figure 8. DBSs of lncRNAs from the same lineage and overlapping TEs from the same lineage either outnumber or are stronger than DBSs of lncRNAs from different lineages and overlapping TEs from different lineages. (A) For DBSs with affinity=>60 (but not =>100), there is a significant difference in affinity. DBSs of PS/SS/HS lncRNAs and overlapping PS/SS TEs are stronger than DBSs of RS/SS lncRNAs and overlapping RS TEs, and DBSs of RS/MS lncRNAs and overlapping RS TEs are stronger than DBSs of PS/SS/HS lncRNAs and overlapping PS/SS TEs (two-sided Mann-Whitney U test). (B) For DBSs with affinity=>80, there is a significant difference in the number. DBSs of PS/SS/HS lncRNAs and overlapping PS/SS TEs outnumber DBSs of RS/MS lncRNAs and overlapping RS TEs, and DBSs of RS/MS lncRNAs and overlapping RS TEs outnumber DBSs of PS/SS/HS lncRNAs and overlapping PS/SS TEs ($P<0.0001$, Chi-Square test).

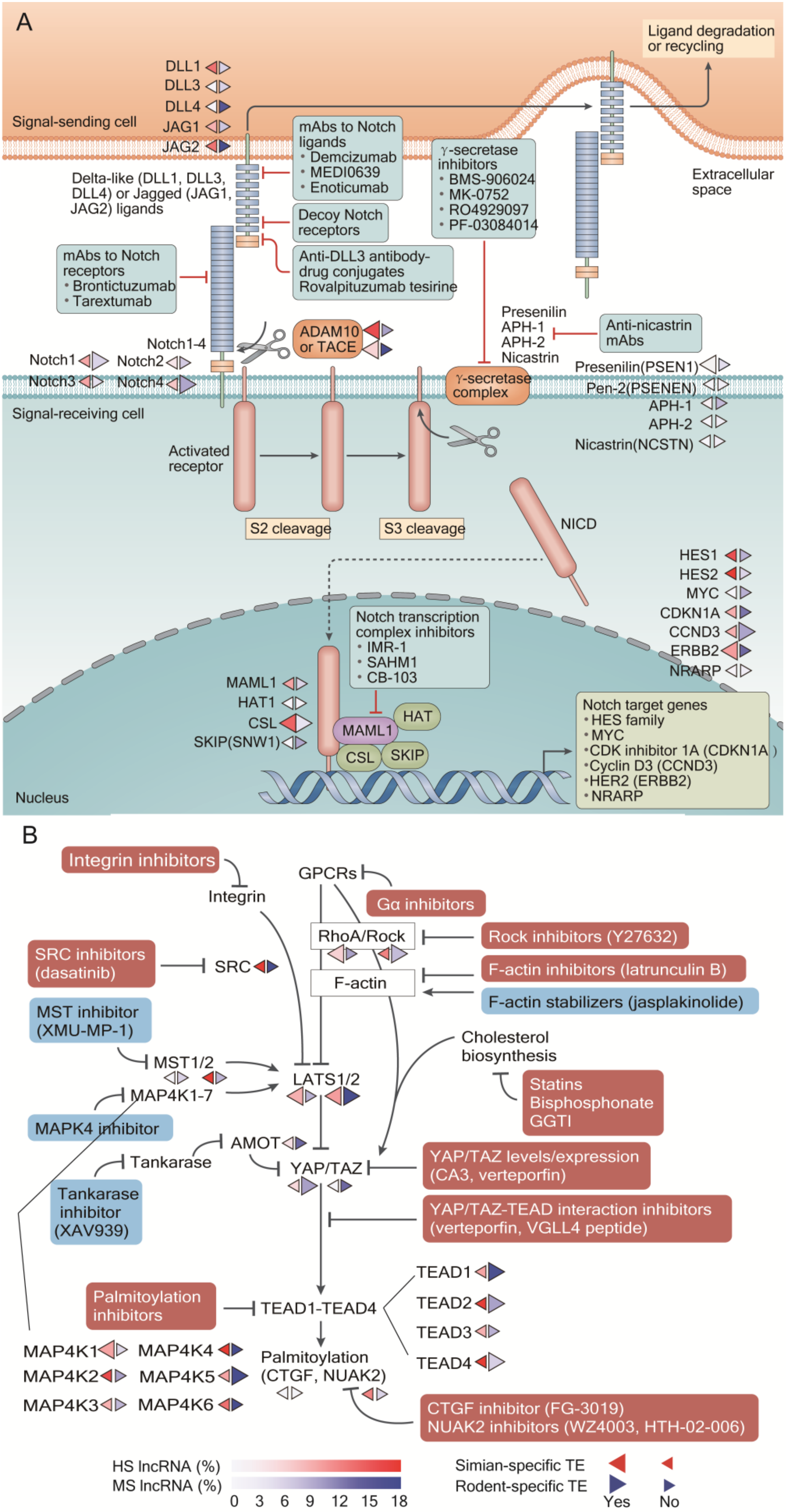
A
Ligand degradation or recycling
Signal-sending cell
DLL1
DLL3
DLL4
JAG1
JAG2
mAbs to Notch ligands
• Demcizumab
• MEDI0639
• Enoticumab
γ-secretase inhibitors
• BMS-906024
• MK-0752
• RO4929097
• PF-03084014
Extracellular space
Delta-like (DLL1, DLL3, DLL4) or Jagged (JAG1, JAG2) ligands
Decoy Notch receptors
Anti-DLL3 antibody-drug conjugates
Rovalpituzumab tesirine
mAbs to Notch receptors
• Brontictuzumab
• Tarextumab
Presenilin
APH-1
APH-2
Nicastrin
Anti-nicastrin mAbs
Notch1-4
ADAM10 or TACE
Notch1
Notch2
Notch3
Notch4
γ-secretase complex
Presenilin(PSEN1)
Pen-2(PSENEN)
APH-1
APH-2
Nicastrin(NCSTN)
Signal-receiving cell
Activated receptor
S2 cleavage
S3 cleavage
NICD
HES1
HES2
MYC
CDKN1A
CCND3
ERBB2
NRARP
Notch transcription complex inhibitors
• IMR-1
• SAHM1
• CB-103
MAML1
HAT1
CSL
SKIP(SNW1)
HAT
MAML1
CSL
SKIP
Notch target genes
• HES family
• MYC
• CDK inhibitor 1A (CDKN1A)
• Cyclin D3 (CCND3)
• HER2 (ERBB2)
• NRARP
Nucleus
B
Integrin inhibitors
Integrin
GPCRs
Gα inhibitors
RhoA/Rock
Rock inhibitors (Y27632)
SRC inhibitors (dasatinib)
SRC
F-actin
F-actin inhibitors (latrunculin B)
F-actin stabilizers (jasplakinolide)
MST inhibitor (XMU-MP-1)
Cholesterol biosynthesis
MST1/2
LATS1/2
Statins
Bisphosphonate
GGTI
MAP4K1-7
MAPK4 inhibitor
AMOT
Tankarase
YAP/TAZ
YAP/TAZ levels/expression (CA3, verteporfin)
Tankarase inhibitor (XAV939)
YAP/TAZ-TEAD interaction inhibitors (verteporfin, VGLL4 peptide)
TEAD1
TEAD2
TEAD3
TEAD4
Palmitoylation inhibitors
TEAD1-TEAD4
MAP4K1
MAP4K4
MAP4K2
MAP4K5
MAP4K3
MAP4K6
Palmitoylation (CTGF, NUAK2)
CTGF inhibitor (FG-3019)
NUAK2 inhibitors (WZ4003, HTH-02-006)
HS lncRNA (%)
MS lncRNA (%)
0 3 6 9 12 15 18
Simian-specific TE
Rodent-specific TE
Yes
No

Supplementary Figure 9. Transcriptional regulation by HS and MS lncRNAs reshapes signaling pathways. The sizes and colors of triangles attached to each gene indicate whether the gene contains simian/rodent TEs and the percentage (from 0% to 18%) of regulatory HS/MS lncRNAs (HS and MS lncRNAs that have DBSs in the gene), respectively. Pathway maps are adapted from the published papers (Clara et al., 2020; Dey et al., 2020). (A) The Notch pathway. (B) The Hippo pathway.

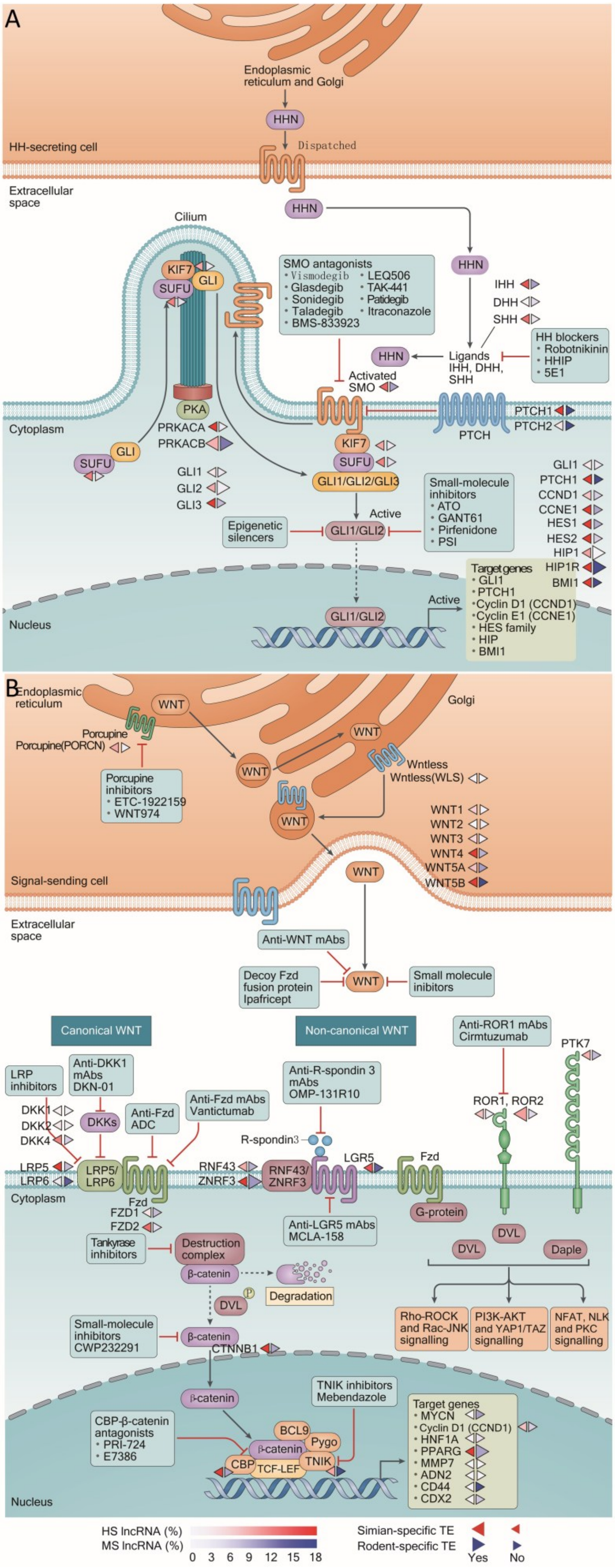
A
Endoplasmic reticulum and Golgi
HHN
Dispatched
HH-secreting cell
Extracellular space
HHN
Cilium
KIF7
GLI
SUFU
SMO antagonists
• Vismodegib • LEQ506
• Glasdegib • TAK-441
• Sonidegib • Patidegib
• Taladegib • Itraconazole
• BMS-833923
HHN
IHH
DHH
SHH
HH blockers
• Robotnikinin
• HHIP
• 5E1
HHN
Ligands IHH, DHH, SHH
Activated SMO
PTCH1
PTCH2
PTCH
PKA
Cytoplasm
PRKACA
PRKACB
GLI
SUFU
KIF7
SUFU
GLI1/GLI2/GLI3
GLI1
GLI2
GLI3
Small-molecule inhibitors
• ATO
• GANT61
• Pirfenidone
• PSI
Active
Epigenetic silencers
GLI1/GLI2
GLI1
PTCH1
CCND1
CCNE1
HES1
HES2
HIP1
HIP1R
BMI1
Target genes
• GLI1
• PTCH1
• Cyclin D1 (CCND1)
• Cyclin E1 (CCNE1)
• HES family
• HIP
• BMI1
Active
Nucleus
GLI1/GLI2
B
Endoplasmic reticulum
WNT
Golgi
Porcupine
Porcupine(PORCN)
WNT
WNT
Wntless
Wntless(WLS)
Porcupine inhibitors
• ETC-1922159
• WNT974
WNT
WNT1
WNT2
WNT3
WNT4
WNT5A
WNT5B
Signal-sending cell
WNT
Extracellular space
Anti-WNT mAbs
Decoy Fzd fusion protein Ipafricept
WNT
Small molecule inibitors
Canonical WNT
Non-canonical WNT
Anti-ROR1 mAbs Cirmtuzumab
PTK7
LRP inhibitors
Anti-DKK1 mAbs DKN-01
Anti-R-spondin 3 mAbs OMP-131R10
DKK1
DKK2
DKK4
DKKs
Anti-Fzd ADC
Anti-Fzd mAbs Vantictumab
ROR1, ROR2
R-spondin3
LGR5
Fzd
LRP5
LRP6
LRP5/LRP6
RNF43
ZNRF3
RNF43/ZNRF3
Cytoplasm
Fzd
FZD1
FZD2
Anti-LGR5 mAbs MCLA-158
G-protein
DVL
DVL
Daple
Tankyrase inhibitors
Destruction complex
β-catenin
Degradation
DVL
Rho-ROCK and Rac-JNK signalling
PI3K-AKT and YAP1/TAZ signalling
NFAT, NLK and PKC signalling
Small-molecule inhibitors CWP232291
β-catenin
CTNNB1
β-catenin
TNIK inhibitors Mebendazole
CBP-β-catenin antagonists
• PRI-724
• E7386
BCL9
β-catenin
Pygo
CBP
TCF-LEF
TNIK
Target genes
• MYCN
• Cyclin D1 (CCND1)
• HNF1A
• PPARG
• MMP7
• ADN2
• CD44
• CDX2
Nucleus
HS lncRNA (%)
MS lncRNA (%)
0 3 6 9 12 15 18
Simian-specific TE
Rodent-specific TE
Yes
No

Supplementary Figure 10. Transcriptional regulation by HS and MS lncRNAs reshapes signaling pathways. The sizes and colors of triangles attached to each gene indicate whether the gene contains simian/rodent TEs and the percentage (from 0% to 18%) of regulatory HS/MS lncRNAs (HS and MS lncRNAs that have DBSs in the gene), respectively. Pathway maps are adapted from the published papers (Clara et al., 2020; Dey et al., 2020). (A) The Hedgehog pathway. (B) The Wnt pathway.

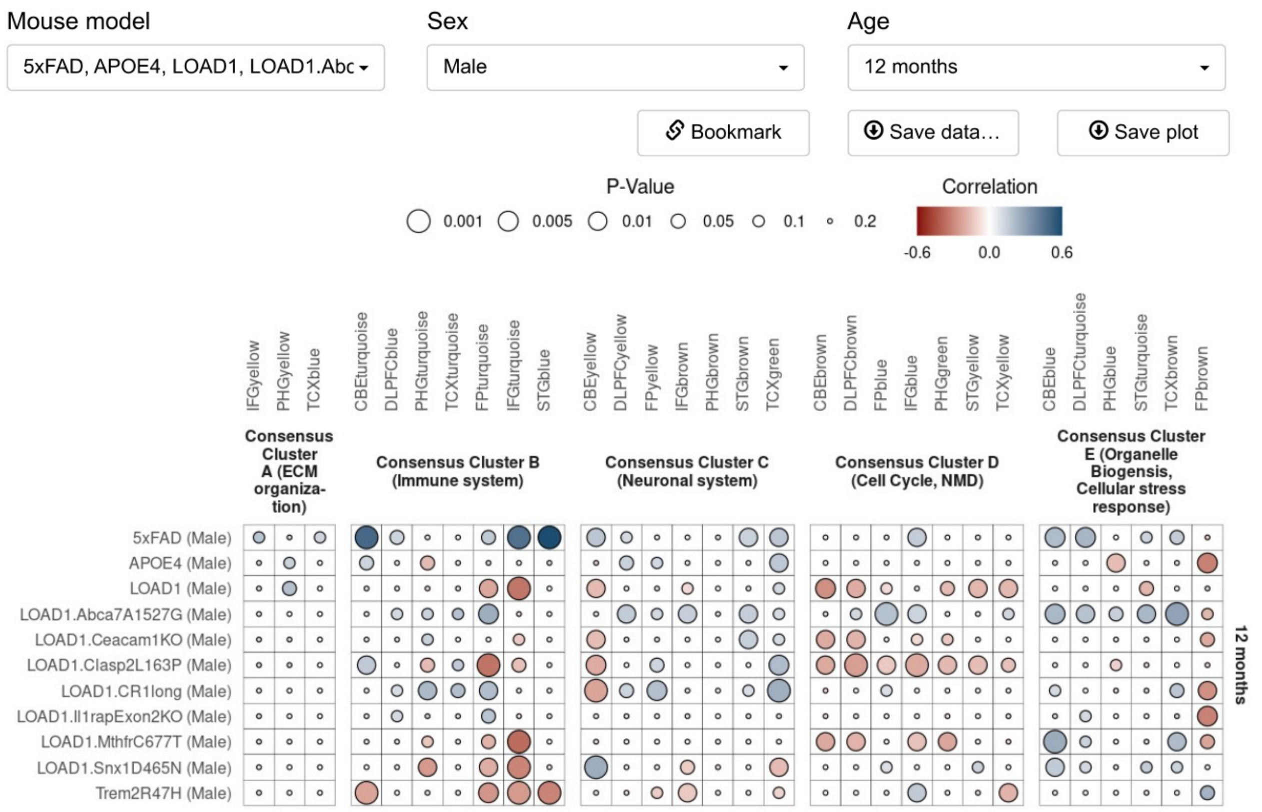


Supplementary Figure 11. This figure is generated by the "Model AD Explore" website (https://sagebio.shinyapps.io/MODEL_AD_Explorer/#) and shows the extent to which gene expression changes in AD mouse models mimic gene expression changes in human AD patients. The left area shows AD mouse models. Gene expression changes in each mouse model are calculated compared to controls; the resulting fold change for each gene is then correlated with the fold change computed based on human AD patients and controls (Preuss et al., 2020). Then, the correlations between mouse models and human patients are mapped to consensus network modules obtained from a harmonized co-expression analysis based on data from three human postmortem brain cohorts (Wan et al., 2020). Blue dots indicate positive correlations (the gene expression changes in mice are similar to changes in humans); red dots indicate negative correlations (the gene expression changes in mice are in the opposite direction of humans); and the sizes of dots indicate P values. Notably, many negative correlations exist between gene expression in mouse models and human patients.

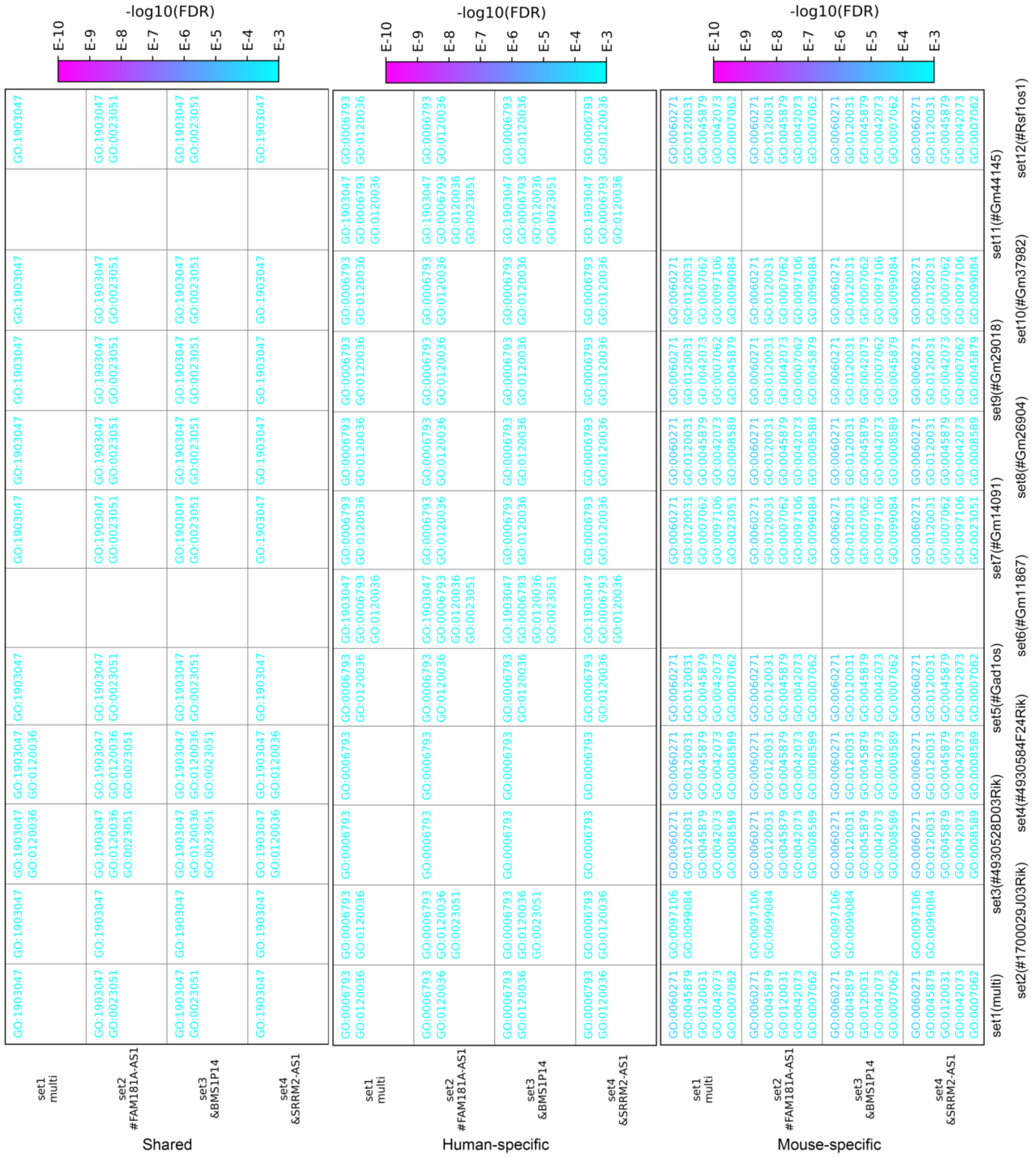


Supplementary Figure 12. Shared, human-specific, and mouse-specific GO BP terms in merged sets in human and mouse RS cells. Each set, with the LS lncRNA (if it has only one initial lncRNA set) or "multi" (if it has multiple initial lncRNA sets) at the bottom and right, indicates a merged set in humans and mice, respectively. GO terms common to most sets and all cells, such as spermatogenesis (GO:0007283), cell cycle (GO:0007049), and system development (GO:0048731), had been filtered out. Colors and numbers in color bars indicate significance levels of enrichment of unfiltered GO terms. Only the top five are displayed if there are >5 sets.

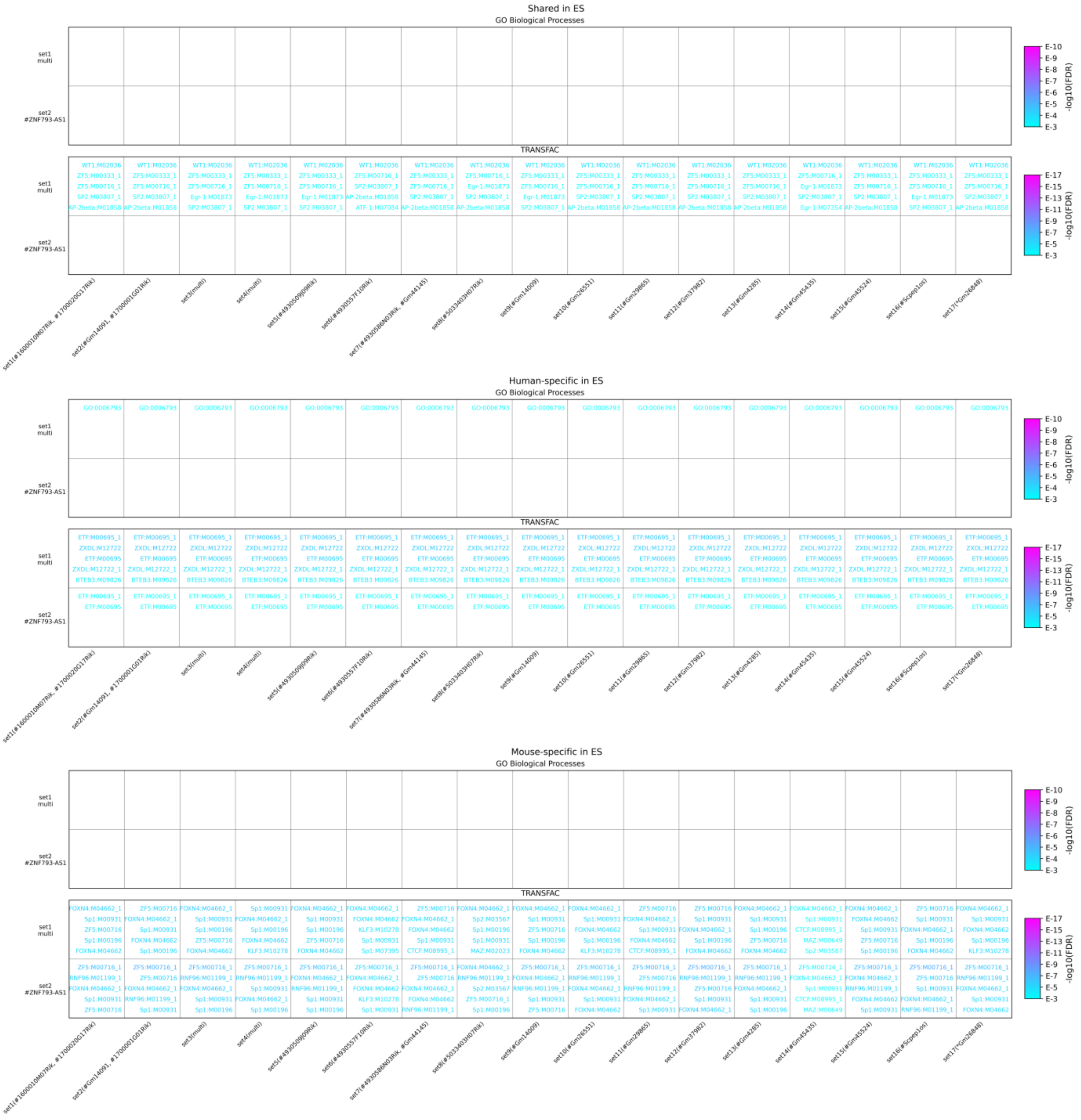


Supplementary Figure 13. Shared, human-specific, and mouse-specific GO BP terms and TRANSFAC terms in merged sets in ES cells in humans and mice. Only the top 5 terms are shown.

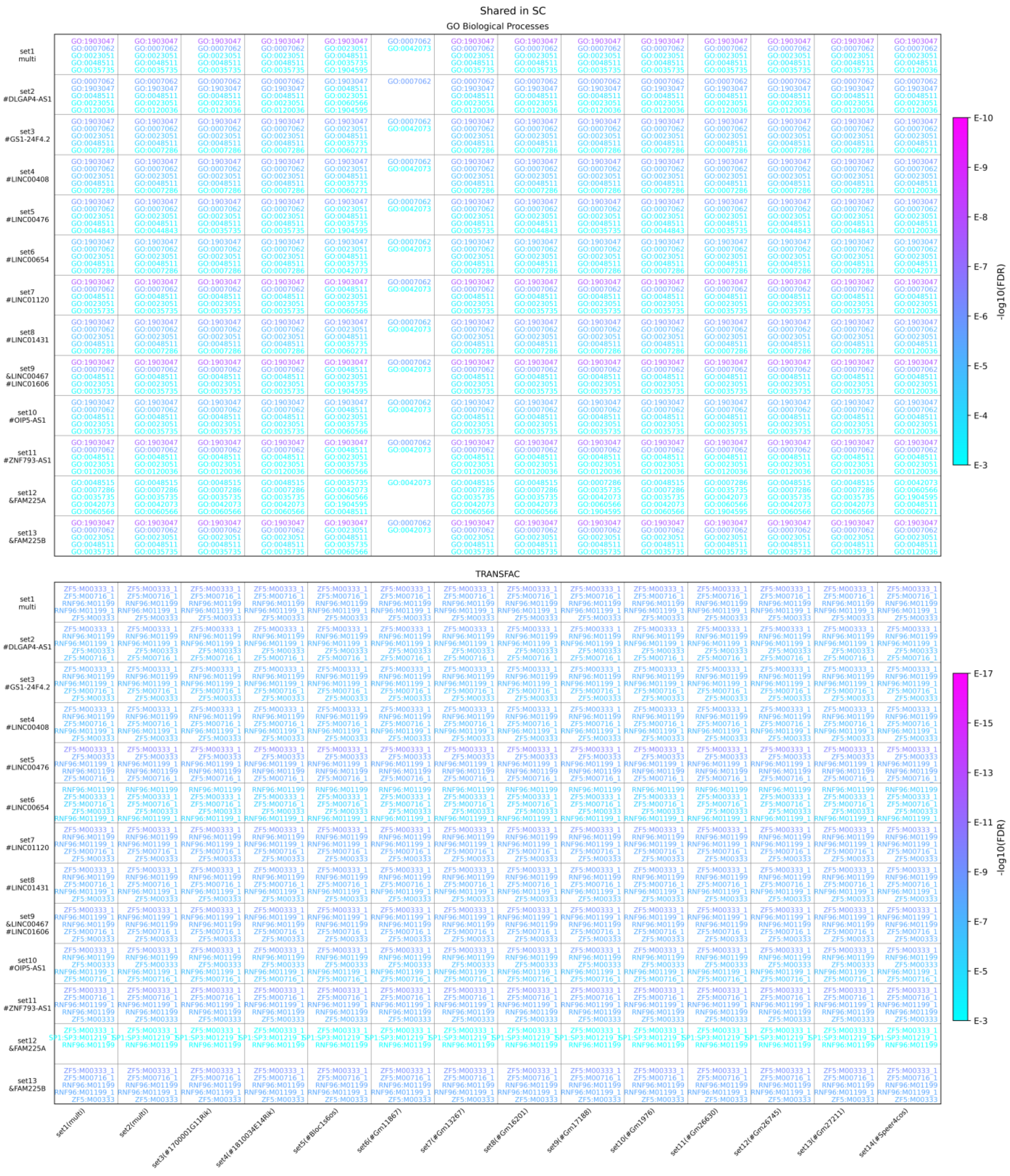


Supplementary Figure 14. (A) The shared GO BP terms and TRANSFAC terms in merged sets in SC cells in humans and mice. Only the top 5 terms are shown.

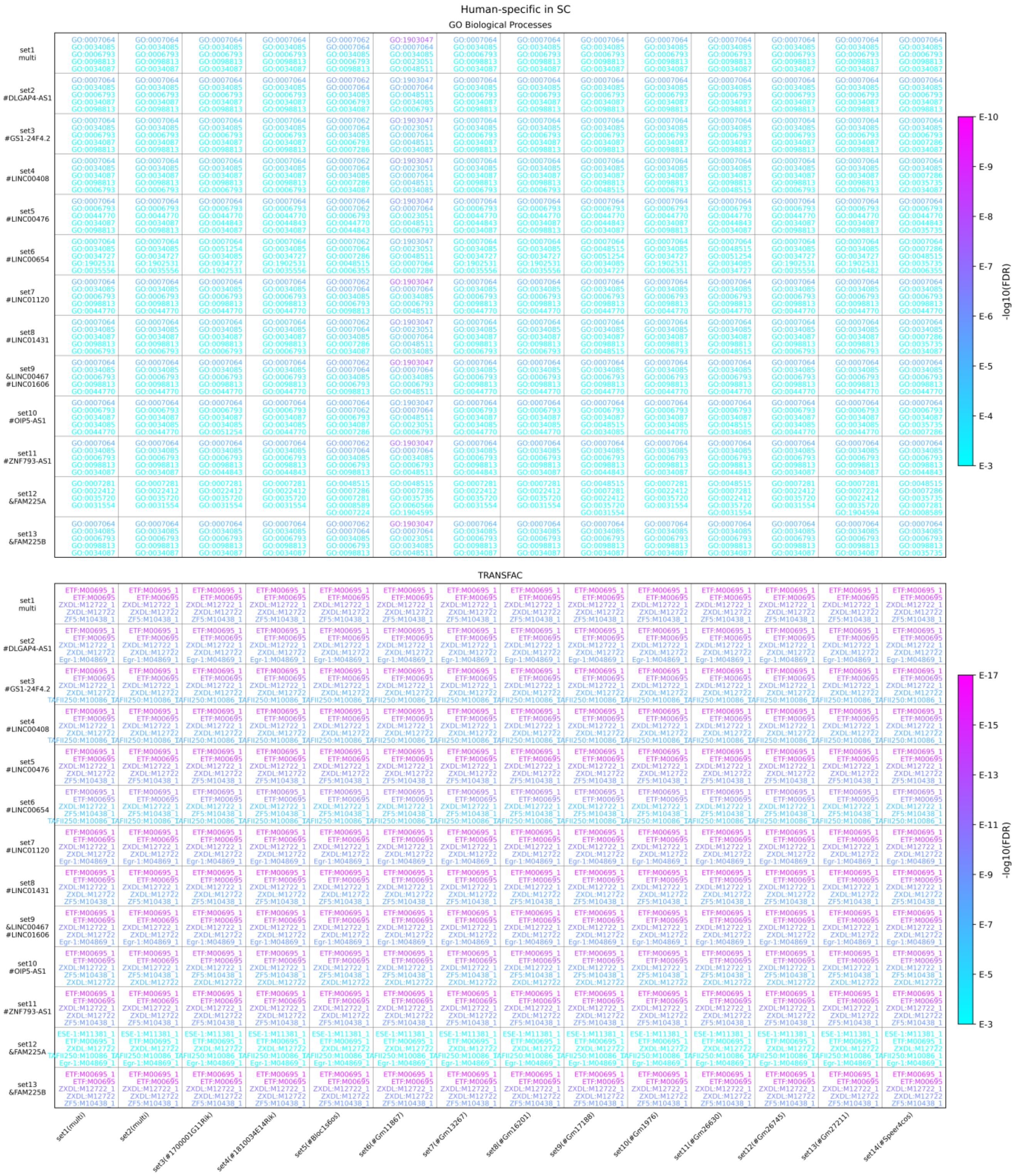


Supplementary Figure 14. (B) The human-specific GO BP terms and TRANSFAC terms in merged sets in SC cells in humans and mice. Only the top 5 terms are shown.

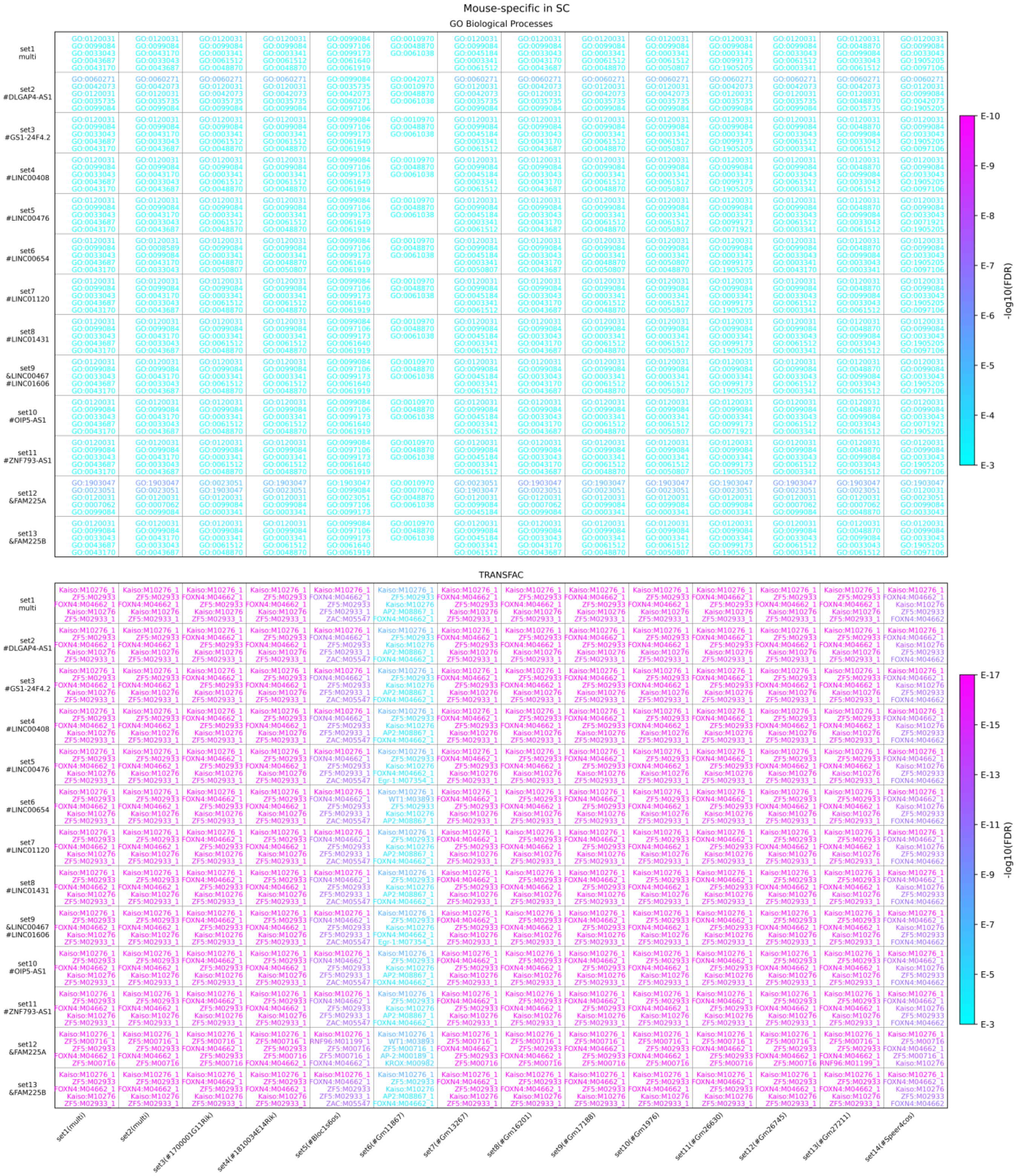


Supplementary Figure 14. (C) The mouse-specific GO BP terms and TRANSFAC terms in merged sets in SC cells in humans and mice. Only the top 5 terms are shown.
